# Identification of Synchronously Active Cells (ISAC) to study the activity of cell assemblies

**DOI:** 10.64898/2026.10.09.758015

**Authors:** Gabriel Makdah, Sidney I. Wiener, Ralitsa Todorova, Marco N. Pompili

**Affiliations:** Institut de Neurosciences des Systémes (INS), INSERM UMR1106, Aix-Marseille Université, 27 Boulevard Jean Moulin, 13005 Marseille, France; Centre Interdisciplinaire de Recherche en Biologie (CIRB) - CNRS UMR 7241 - INSERM U1050, Collége de France, Université PSL, 11 place Marcelin Berthelot, 75005 Paris, France

## Abstract

Cell assemblies–groups of neurons that fire synchronously–are central to neuronal computation. Technological advances now permit the simultaneous recording of large neuronal populations; however, many existing cell assembly detection algorithms rely on simplified measures such as pairwise correlations, failing to capture higher-order synchrony and thereby missing a rich repertoire of assemblies present in the data. To address these limitations, we developed Identification of Synchronously Active Cells (ISAC), which performs a rigorous, iterative search across all recorded units, leveraging the full dimensionality of the dataset to detect groups that fire together above chance. Using simulated recordings with ground-truth assemblies of increasing complexity, we demonstrate that ISAC considerably outperforms currently used algorithms in identifying synchronously active cells. Finally, we show that applying ISAC to experimental recordings uncovers novel features of assembly formation and dynamics. The power and sensitivity of ISAC can thus enable insights that remain inaccessible with existing methods.

**SIGNIFICANCE STATEMENT:** Neurons rarely act in isolation. Instead, groups of neurons can become active together, forming “cell assemblies” thought to represent and process information in the brain. Identifying which neurons belong to these assemblies is therefore important for understanding how neuronal populations encode information. However, commonly used methods can incorrectly merge distinct groups of neurons or identify neurons that do not consistently participate in synchronous activity. Here, we introduce Identification of Synchronously Active Cells (ISAC), a method designed to systematically identify groups of neurons that fire together more often than expected by chance. ISAC directly examines the timing of neuronal activity and applies statistical criteria to determine both whether individual neurons belong to a group and whether the group behaves as a coherent assembly. Across extensive simulations, including conditions reproducing the complex activity patterns observed in experimental recordings, ISAC identified cell assemblies more accurately than commonly used approaches. Applications to real recordings further illustrate how ISAC can reveal the formation, composition, spatial coding, and reactivation of neuronal assemblies. ISAC provides a framework for studying how coordinated groups of neurons organize and process information across brain states and behaviors.

## INTRODUCTION

Understanding how neuronal activity leads to cognition and behavior remains a central challenge in neuroscience. A prevailing hypothesis posits that these complex functions emerge from the synchronous activity of specific neuronal ensembles, cell assemblies, which act as fundamental units of neural computation (Buzśaki 2010; Eichenbaum 2018; Hebb 1949). Methodological advances over the last two decades have enabled the simultaneous recording of large numbers of neurons across multiple brain regions (Berényi et al. 2013; Brinkmann et al. 2009; Buzśaki 2004; Einevoll et al. 2012; Jun et al. 2017; Khodagholy et al. 2015; Steinmetz et al. 2021; Wang et al. 2024). Such large-scale recordings present the opportunity to investigate the cell assembly dynamics underlying a wide range of cognitive phenomena. Indeed, cell assemblies have been shown to be involved in perception (Filipchuk et al. 2022; Luczak et al. 2009), memory (Benchenane et al. 2010; El-Gaby et al. 2021; Liu et al. 2023; Miyawaki and Mizuseki 2022; Panichello et al. 2024; Peyrache et al. 2009; Pompili et al. 2026, 2025b; Ramanathan et al. 2015; Robinson et al. 2025; Todorova and Zugaro 2019), and executive and motor functions (Carmena et al. 2003; Dejean et al. 2016; Fujisawa et al. 2008; Georgopoulos et al. 1986; Oberto et al. 2022; Russo et al. 2024; Wessberg et al. 2000). Yet, despite these technological advancements, the existing tools for cell assembly detection (Billeh et al. 2014; Humphries 2011; Lopes-dos-Santos et al. 2013; Maboudi et al. 2018; Mackevicius et al. 2019; Peyrache et al. 2010; Russo and Durstewitz 2017) have several theoretical and practical limitations, some of which have been highlighted in previous work (Deolindo et al. 2014; Makdah et al. 2025; Sakurai et al. 2018).

Detecting groups of synchronously active cells is a computationally demanding task and many of the established algorithms use clever shortcuts such as pairwise correlation measures to infer cell assembly membership (Diana et al. 2019; Humphries 2011; Lopes-dos-Santos et al. 2013; Peyrache et al. 2010; Tavoni et al. 2016). However, groups of pairwise-correlated cells do not necessarily inform recurrent patterns of co-activation, since cells could fire together in pairs without the entire group ever firing all together, especially given that neurons have been shown to participate in overlapping sets of assemblies (Alme et al. 2014; El-Gaby et al. 2021; Leutgeb et al. 2004; Muller and Kubie 1987; O’Keefe and Conway 1978; Pompili et al. 2026, 2025b). For example, consider the following simplified case: suppose cells A, B, and C each co-activate with different sets of other cells, forming assemblies that never include all three together (**Figure 1A**). Even though A, B, and C never fire simultaneously, each pair (AB, AC, BC) can show strong correlations, which could mislead an algorithm relying on pairwise correlations to group them as a single assembly (**Figure 1B**).

**Figure 1:**
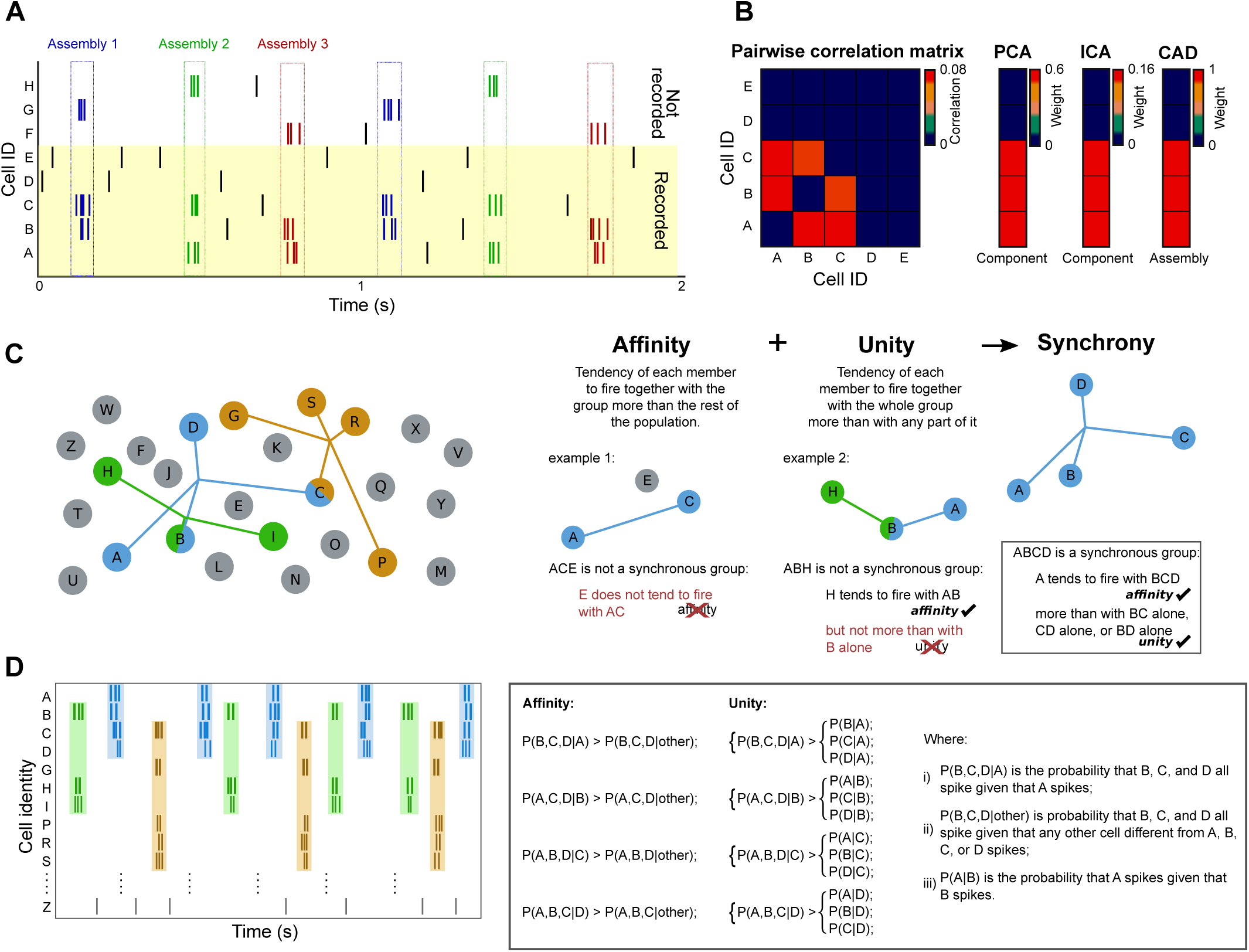
Defining groups of Synchronously Active Cells (SAC assemblies). (A) Raster plot of a simulated group of neurons containing three cell assemblies. Nonrecorded (top) and recorded (bottom, shaded) neurons are both shown. Note how recorded neurons (cells A to E) participate in multiple overlapping assemblies. (B) Pairwise correlation matrix (left) of recorded neurons in (A) and the putative cell assemblies detected using the methods PCA, ICA and CAD (right). PCA: principal component analysis (Peyrache et al. 2010); ICA: Independent Component Analysis (Lopes-dos-Santos et al. 2013); CAD: Cell Assembly Detection (Russo and Durstewitz 2017). Note how A, B, and C are detected as an assembly even though they never fire together in panel (A). (C) Schematic representation of the synchrony relationships within a simulated group of neurons A to Z (left). Radial spokes extend to members of color-coded assemblies. Each color-coded group comprises neurons that are members of a SAC assembly. Note how certain neurons participate in multiple SAC assemblies. Right: illustrations of the two statistical criteria defining a SAC assembly. Affinity ensures a neuron fires significantly more with the candidate SAC assembly than the general population does. Unity ensures a neuron fires with the collective group significantly more than with any subset of it. (D) Left: Raster plot of an example simulation of the neurons represented in (C). Colored boxes represent activations of synchronously active cells. Right: Mathematical formulation of the Affinity and Unity criteria for group ABCD.

Addressing these limitations requires moving beyond pairwise measures and low-dimensional projections to methods that account for the high-dimensional nature of neuronal population activity. Given that computational capabilities of devices commonly available for data analysis have greatly advanced since the earliest techniques developed to study cell assemblies, computational shortcuts may no longer be necessary for cell assembly detection, and it should now be possible to comprehensively detect groups of neurons active synchronously. However, to the best of our knowledge, no existing algorithms for the detection of cell assemblies verify that all cells which compose the cell assembly are active together above random chance. As such, while the patterns detected by current algorithms do represent network relationships present in the data, they do not necessarily represent groups of cells that fire together.

Dimensionality reduction and manifold analysis are valuable descriptors of collective dynamics. Low-dimensional manifolds summarize population activity, but they are not designed to identify groups of neurons that fire together as a unit and, by construction, they compress activity into a small number of factors, whereas in principle cell assemblies—defined as combinations of co-active neurons—can be far more numerous than the recorded cells themselves.

To address these issues, enable the direct and comprehensive detection of synchronous patterns, and exploit the full dimensionality of high-density recordings we developed Identification of Synchronously Active Cells (ISAC). Rather than use pairwise correlations as a proxy measure of cellular co-activations, ISAC directly searches for and detects groups of cells firing together above random chance levels. The resulting patterns are composed only of members that are recurrently active together, ensuring that individual groups are not erroneous amalgams of distinct assemblies. Furthermore, ISAC conducts an iterative search for all possible combinations of neurons within a recording, ensuring the detection of virtually all significantly active groups of synchronously active cells present.

Here, we first define statistical criteria to test if groups of neurons fire together more than by chance. Then, we test leading assembly detection methods against these criteria on experimentally recorded data, showing that many of the detected putative cell assemblies fail to fire together. We also used a wide array of artificially generated benchmark datasets, where existing algorithms do not reliably identify the ground truth groups of synchronous neurons. To solve these problems, we present ISAC, and demonstrate its superior accuracy in identifying groups of synchronous ensembles in simulated ground truth data compared to currently leading algorithms. Finally, in experimentally recorded datasets, we show that ISAC is a powerful tool for the detection of cell assemblies in neural recordings and opens the door for novel analyses of cell assemblies.

## RESULTS

### Cell assemblies as groups of synchronous units

The Hebbian concept of a cell assembly, and its contemporary interpretations (e.g., Buzśaki (2010)), emphasize the synchronous co-activation of its constituent neurons as fundamental to its computational role. In practice it is very unlikely to record the activity of all the members of a cell assembly in vertebrate brains (it is possible in some invertebrate preparations) as only a subsample of all the neurons of the brain are recorded at a given moment. Rather, groups of synchronously active cells among a set of recorded units are quite likely parts of larger unrecorded (and therefore unobservable) assemblies. Yet synchronous firing of all members is not a formal requirement of currently employed cell assembly detection algorithms [(Billeh et al. 2014; Humphries 2011; Lopes-dos-Santos et al. 2013; Maboudi et al. 2018; Mackevicius et al. 2019; Peyrache et al. 2010; Russo and Durstewitz 2017) see Makdah et al. (2025) for a review]. However, there is no established consensus on formal criteria for defining when a group of neurons should be considered synchronously active. Here we propose two criteria to establish our working definition of a cell assembly, and we will refer to groups fulfilling the criteria as Synchronously Active Cell assemblies (SAC assemblies) to avoid confusion with other methods. Within this framework, a SAC assembly is characterized by two properties: *affinity* between all members and *unity* of the group (**Figure 1C-D**).

Affinity measures how strongly a neuron is coupled to the group as a whole: each putative member must fire with the rest of the group more often than the general population does. This prevents non-stationary changes in population activity such as fluctuations in global firing rate from being misclassified as SAC assemblies. Unity describes the preference of each member to fire with the collective activation of the full set of SAC assembly neurons rather than with any subset. For example, for the group ABCD to satisfy the unity criterion, neuron A must fire significantly more with the collective activation of BCD than with any smaller combination such as BC, CD, or BD. This ensures that apparent higher-order synchrony cannot be explained by recurring lower-order co-activation patterns among subsets of neurons, so distinct SAC assemblies sharing overlapping neurons (e.g., ABCD and CDEF) are not merged due to shared low-order correlations. Thus, affinity and unity are complementary criteria: affinity ensures that all members are specifically and consistently engaged with the group, while unity ensures that the SAC assembly functions as a coherent, indivisible co-firing ensemble. Importantly, spike trains are not discretized into temporal bins. Instead, synchrony is evaluated directly from the continuous spike times: spikes from different neurons are considered part of the same activation event when they occur within a given Δ*t*. This avoids artefacts associated with assigning spikes to fixed, non-overlapping temporal bins. Together, these criteria ensure that SAC assemblies reflect coordinated activity that cannot be explained solely by global population fluctuations, independent firing, or partial co-activations among subsets of neurons.

To assess whether putative cell assemblies identified by currently most used algorithms fulfill above criteria, we implemented them (Billeh et al. 2014; Humphries 2011; Lopes-dos-Santos et al. 2013; Maboudi et al. 2018; Mackevicius et al. 2019; Peyrache et al. 2010; Russo and Durstewitz 2017) on prefrontal cortex recordings (Peyrache et al. 2018) with well documented cell assembly activity (Benchenane et al. 2010; Maggi and Humphries 2022; Maggi et al. 2018; Peyrache et al. 2009; Singh et al. 2019). We then evaluated the synchrony of groups composed of 3 or more neurons identified by each of these algorithms. Because ground truth is unknown in experimental recordings, this analysis does not evaluate detection accuracy, but rather tests whether the identified groups satisfy the proposed criteria for synchronous co-activation. Surprisingly, the vast majority (mean 94.3%) of the identified groups failed to pass the affinity and unity criteria, indicating they may not represent SAC assemblies (**Figure 2A**, **Supp. Fig. 1**). This high failure rate of these algorithms led us to question whether SAC assemblies might simply be rare within these datasets. Nonetheless, testing all possible three-cell combinations of units from these recordings, we found 1537 groups (17.9 per session on average) that passed our criteria to be considered SAC assemblies (**Supp. Fig. 2A**, example in **Fig. 2B-D**). Some of these were contained in the assemblies detected by existing methods or correlated with them (**Supp. Fig. 2B**), suggesting that currently employed algorithms are sensitive to co-firing relationships between units but may merge multiple SAC assemblies in the same group. Indeed, we tested this in simulated recordings with overlapping SAC assemblies (i.e. sharing members), and these methods tended to combine ground truth assemblies into *mixed groups* (**Figure 2E**; **Supp. Fig. 3**), although *non-synchronous groups* (i.e. groups of cells not participating together in any ground truth assembly) were also present. This is problematic because mixing SAC assemblies can make it hard to tell whether any one of them carries a signal of interest, because their distinct signals can become diluted or canceled when combined into a single group. Taken together, these findings show that detecting individual assemblies of synchronously active cells may require a different approach.

**Figure 2:**
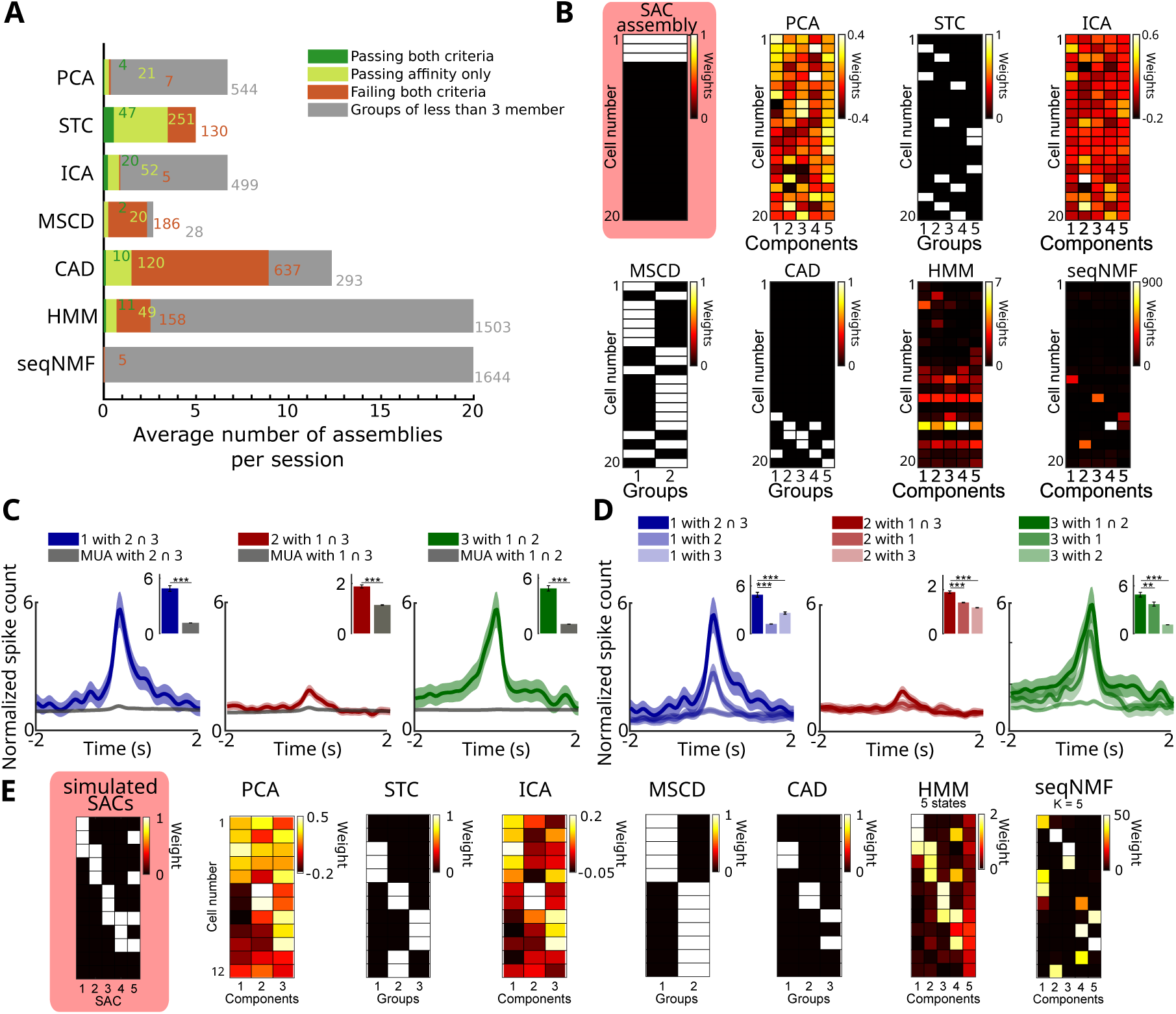
Currently available methods fail to identify SAC assemblies. (**A**) Cumulative count of putative cell assemblies detected by seven established methods in a published set of rat prefrontal cortex recordings, respecting the statistical criteria to define SAC assemblies. The numbers on the bars indicate the total count of assemblies identified by each method across all sessions. Note that affinity and unity are not defined for groups of fewer than 3 members, so only groups of 3 or more members could be tested. (**B**) Putative cell assemblies detected by seven established methods on one example recording of 20 neurons in the rat prefrontal cortex. On the left, highlighted in red, is an example of a synchronously active cell assembly (SAC assembly, adhering to the affinity and unity criteria) present in the data that was not detected by any of the other methods. Each matrix displays neurons (rows) and their participation in detected assemblies (columns). This example SAC assembly is composed of neurons 1, 2, and 3. Methods like PCA, ICA, HMM, and seqNMF produce weighted components (color-coded), while STC, MSCD, and CAD produce groups with all or none participation of neurons (white indicates membership). Note that none of the methods detected this SAC assembly. (**C**) The affinity criterion applied to the example assembly in (B). Each plot compares the normalized peri-event time histogram of the firing of one neuron (e.g. blue line for neuron 1) centered on moments when the other two members are active (e.g. neurons 2 and 3 for the first plot) versus the normalized peri-event time histogram of the entire neural population (non-member population activity; yellow line) centered on those same moments (when the other two members are active). Insets compare the average spike count between-200 and +200 ms (\*\*\**p <* 0.001; Wilcoxon rank-sum test). (**D**) The unity criterion applied to the same SAC assembly. Each plot shows the normalized peri-event time histogram of a given neuron’s firing rate (e.g., left plot is for neuron 1) during the simultaneous activation of all other SAC assembly members (blue line), versus activation of partial subsets (red and green lines). Insets same as (C) (\*\*\**p <* 0.001, \*\**p <* 0.01; Wilcoxon rank-sum test). (**E**) Results of currently used methods in detecting cell assemblies in a small simulated recording of 12 neurons participating in five simulated SAC assemblies (left and highlighted in red, see Methods for details). Same format as (B). HMM and seqNMF require the user to pre-define the number of components to be find, which in this case was set to 5. CAD: Cell Assembly Detection; HMM: Hidden Markov Models; ICA: Independent Component Analysis; MSCD: Markov Stability for Community Detection; PCA: principal component analysis; seqNMF: sequential non-negative matrix factorization; STC: Spike Train Communities.

### A novel approach: Identification of Synchronously Active Cells (ISAC)

To overcome these limitations and methodically detect the cell assemblies that pass the SAC criteria, free from amalgams of asynchronous groups of cells, we developed ISAC. ISAC employs a systematic, iterative approach to identify all neuronal groups that satisfy the affinity and unity criteria defined above (see **Figure 3** for a schematic overview). Note that to avoid ambiguity, in this manuscript we distinguish between an assembly’s definition and its method of detection: a “SAC assembly” is any group satisfying the SAC criteria, regardless of the detection method, whereas an “ISAC assembly” is a SAC assembly identified by the ISAC algorithm.

**Figure 3:**
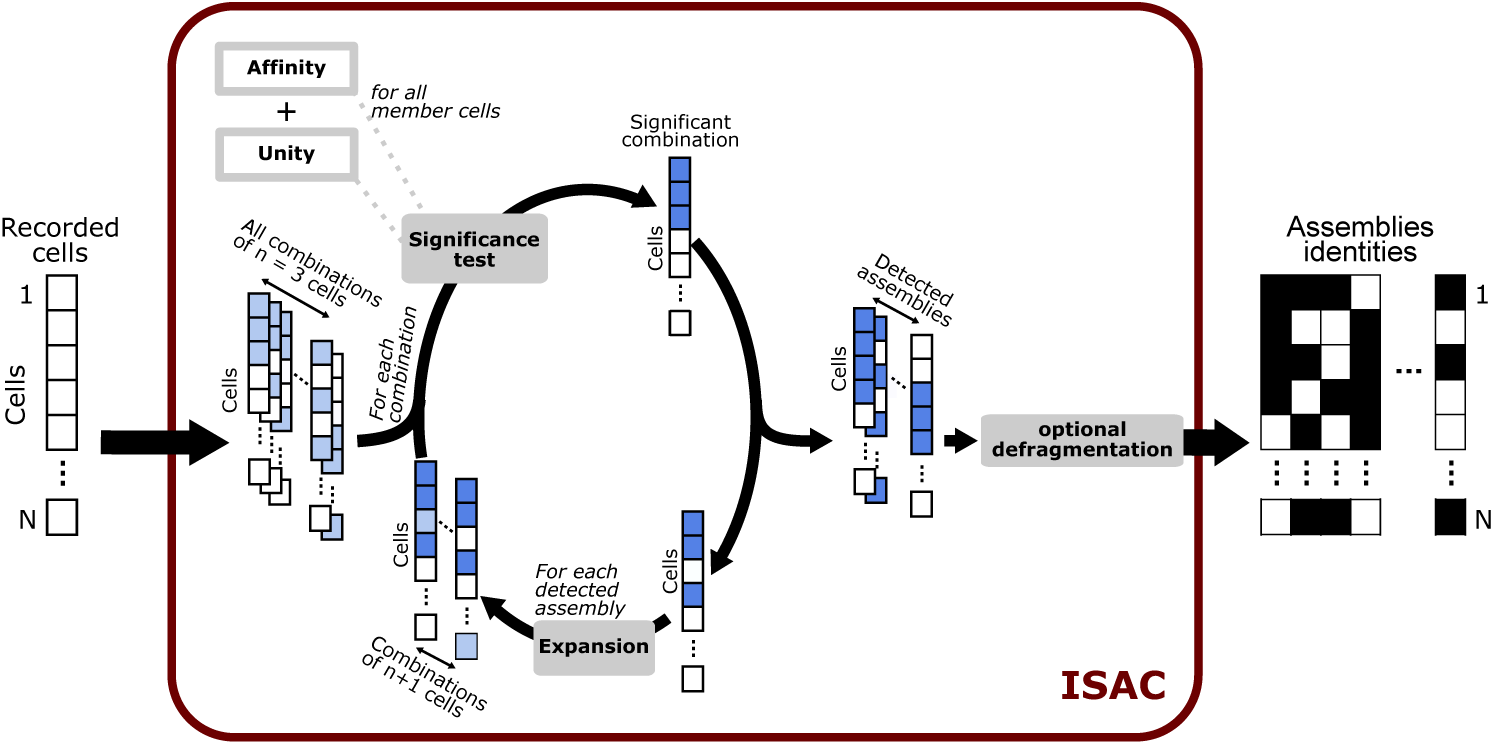
ISAC algorithm for identifying SAC assemblies. The algorithm begins by considering every possible combination of three cells as a candidate SAC assembly (left). A SAC assembly is selected if all its members satisfy the criteria for affinity and unity. Following the significance test step, ISAC performs an expansion step in an iterative manner. For each previously detected SAC assembly, the expansion step consists in finding one additional neuron that may be added to the SAC assembly. Each such 4-neuron combination is evaluated against the affinity and unity criteria. If these statistical criteria are met, the 4-cell combination is considered as a significant 4-member SAC assembly and the cycle continues, progressively appending more neurons until no significant SAC assemblies are detected in a cycle. Finally, an optional defragmentation step consolidates smaller assemblies into a larger SAC assembly that satisfy the affinity criterion (see Methods).

The first step of the algorithm (**Figure 3A**) is to test all possible 3-cell combinations as candidate assemblies. Those triplets that pass the two criteria are retained and serve as a basis for expansion in the next step. For example, if ABC passed, all 4-cell SAC assemblies containing ABC (e.g. ABCD, ABCE, ABCF, etc…) would be the candidate SAC assemblies to be tested according to the criteria in the next iteration. This procedure is repeated iteratively and thus cells are progressively added to SAC assemblies until a cycle in which no SAC assemblies fulfilling affinity and unity are found. Each candidate group is evaluated independently, and every group satisfying the affinity and unity criteria is retained as a distinct SAC assembly. Consequently, ISAC does not attempt to partition neurons into mutually exclusive assemblies. If two overlapping groups (e.g., ABCD and ABXY) both satisfy the statistical criteria, both are reported independently. There is no limit to the number of assemblies in which a given neuron may participate. The algorithm outputs all the SAC assemblies which were retained on previous cycles. ISAC systematically applies these steps to ensure the detection of the SAC assemblies present within a dataset. The only parameters required by the user are the alpha level (Z, set to 2.57 here, corresponding to an estimated false-positive rate of 0.2%; **Supp. Fig. 4**) and the timescale of synchrony (Δ*t*), which was set to 30 ms throughout the present study (unless indicated otherwise), consistent with previous work suggesting this timescale as an upper bound for downstream neuronal integration of synchronous inputs (Pompili et al. 2025b). Finally, an optional defragmentation step with relaxed criteria combines highly overlapping SAC assemblies into a larger assembly.

**Figure 4:**
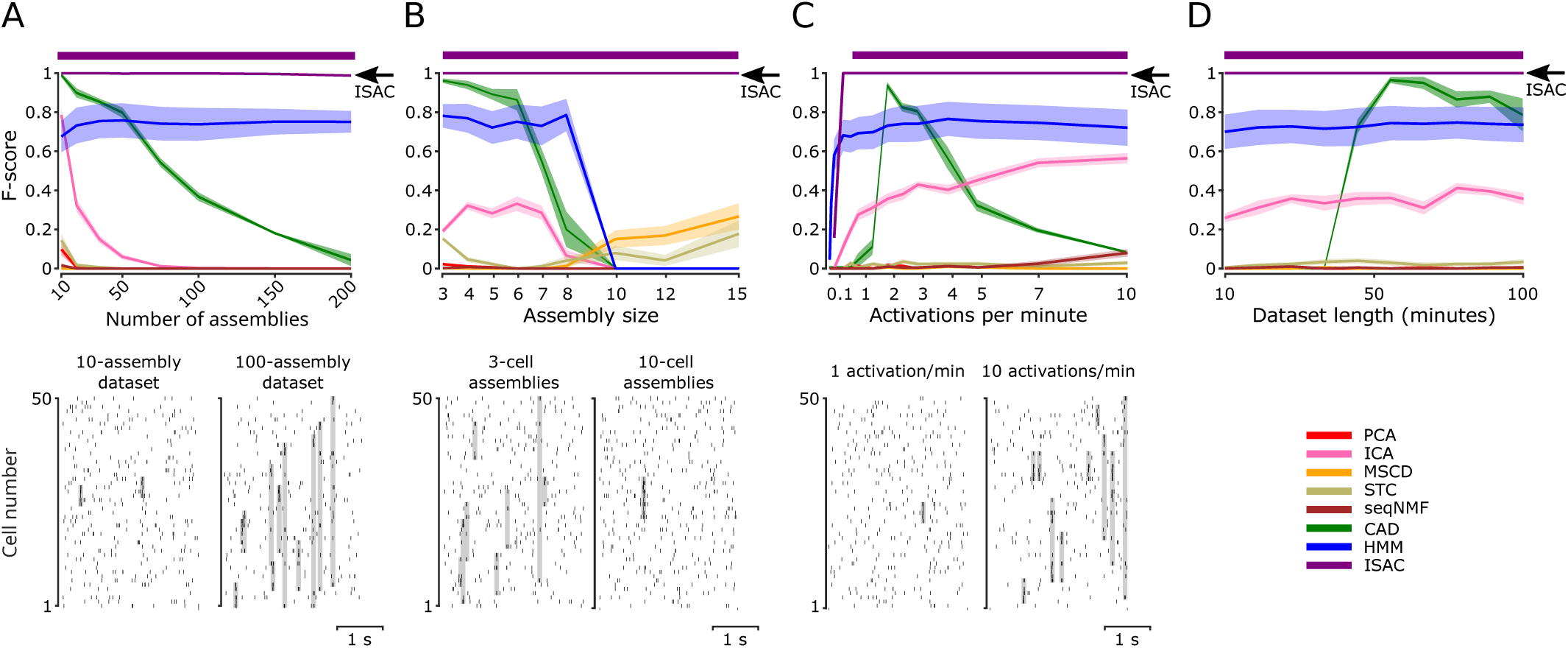
Comparative performances of currently used algorithms and ISAC on simulated recordings containing simulated SAC assemblies embedded in Poisson point process background activity. (A) Top: Average F-score as a function of the number of SAC assemblies. Shading of the curves represent the standard error of the mean (s.e.m.). The thick horizontal line above indicates the data points where the F-score of ISAC is significantly higher than every other method (*p <* 0.05). Bottom: examples of 3 seconds of two simulated recordings with assembly activations highlighted in gray. (B) Same as (A), but varying the number of neurons per assembly. (C) Same as (A), but varying the number of assembly activations per minute. (D) Same as (A), but varying the recording length.

The algorithm described here can thus identify the cell assemblies (i.e. defining which cells are the members of a given group) that satisfy the criteria present in the data. Once the groups are defined, their activity over time can be quantified using various approaches, a problem that is distinct from cell assembly detection. For example, assembly activity can be represented as a simple on/off state or as a continuous activation strength based on the number of active member cells (**Supp. Fig. 5**).

**Figure 5:**
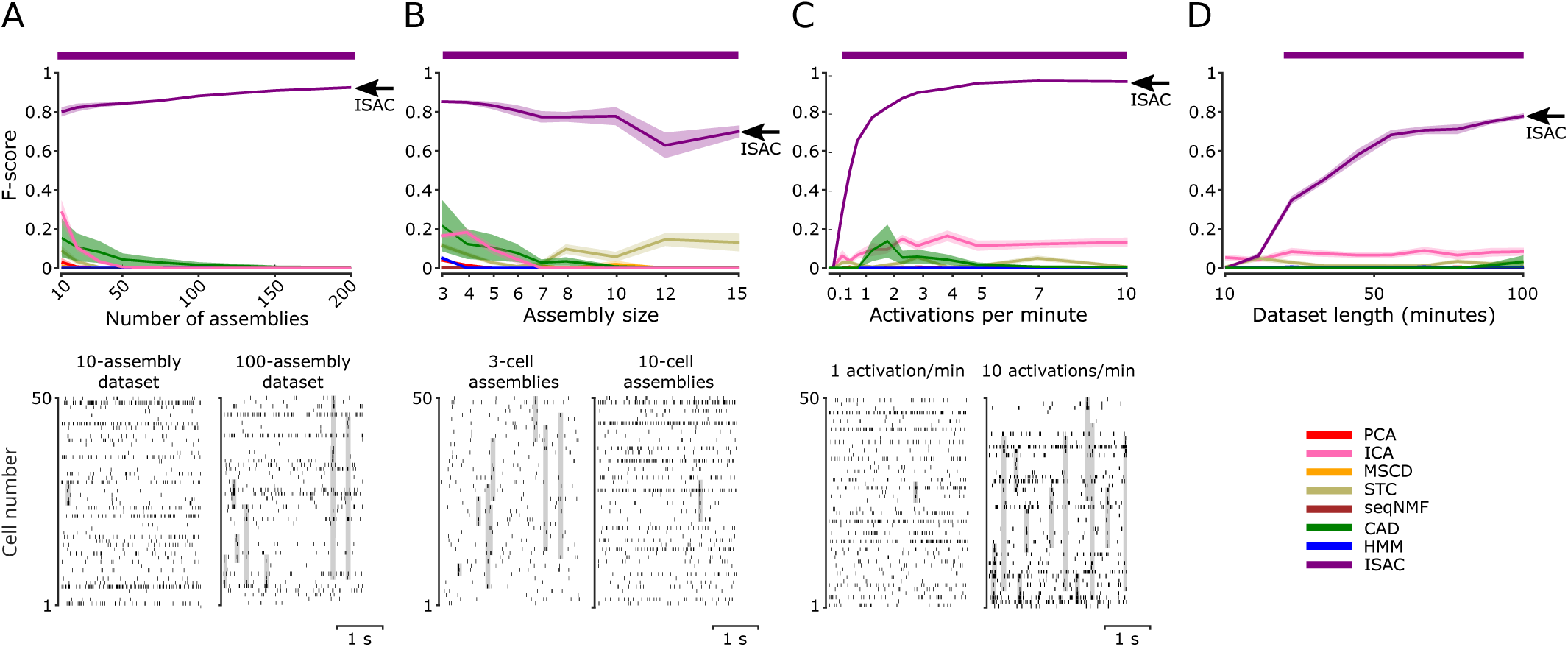
Comparative performances of currently used algorithms and ISAC on simulated recordings containing simulated SAC assemblies embedded in background activity of shuffled prefrontal cortex recordings. Same as Figure 4 but using background activity from published experimental recordings of prefrontal cortical neurons with shuffled neuron identities assigned to individual spikes, preserving spiking statistics (see Methods).

### Algorithm performance on simulated data

To evaluate the performance of ISAC in identifying cell assemblies satisfying the SAC criteria and to compare it to existing algorithms, we generated simulated datasets containing a known ground truth of activation of assemblies embedded within background activity modeled as homogeneous Poisson point processes, representing random uncorrelated firing. These simulations provide an idealized benchmark in which background activity is stationary and homogeneous, allowing algorithm performance to be evaluated under controlled conditions. Fundamental characteristics of cell assemblies in vivo, such as their typical size (the number of neurons per assembly) or their prevalence (the number of assemblies in a given recording) are currently unknown. We therefore varied the number and size of the ground truth assemblies, assessing the scalability of the methods on datasets with overlapping assemblies. While the performance of other algorithms degraded with larger or more numerous assemblies, ISAC performance was consistently high (**Figure 4A-B**, **Supp. Fig. 6A-B**). On the other hand, varying the assembly activation rate and the duration of the simulated recording allowed us to gauge the algorithms’ sensitivity for detecting ‘weaker’ assemblies or performing in datasets of duration which may need to be limited by the experimental paradigm. Here too, ISAC outperformed all other algorithms (**Figure 4C-D** and **Supp. Fig. 6C-D**).

**Figure 6:**
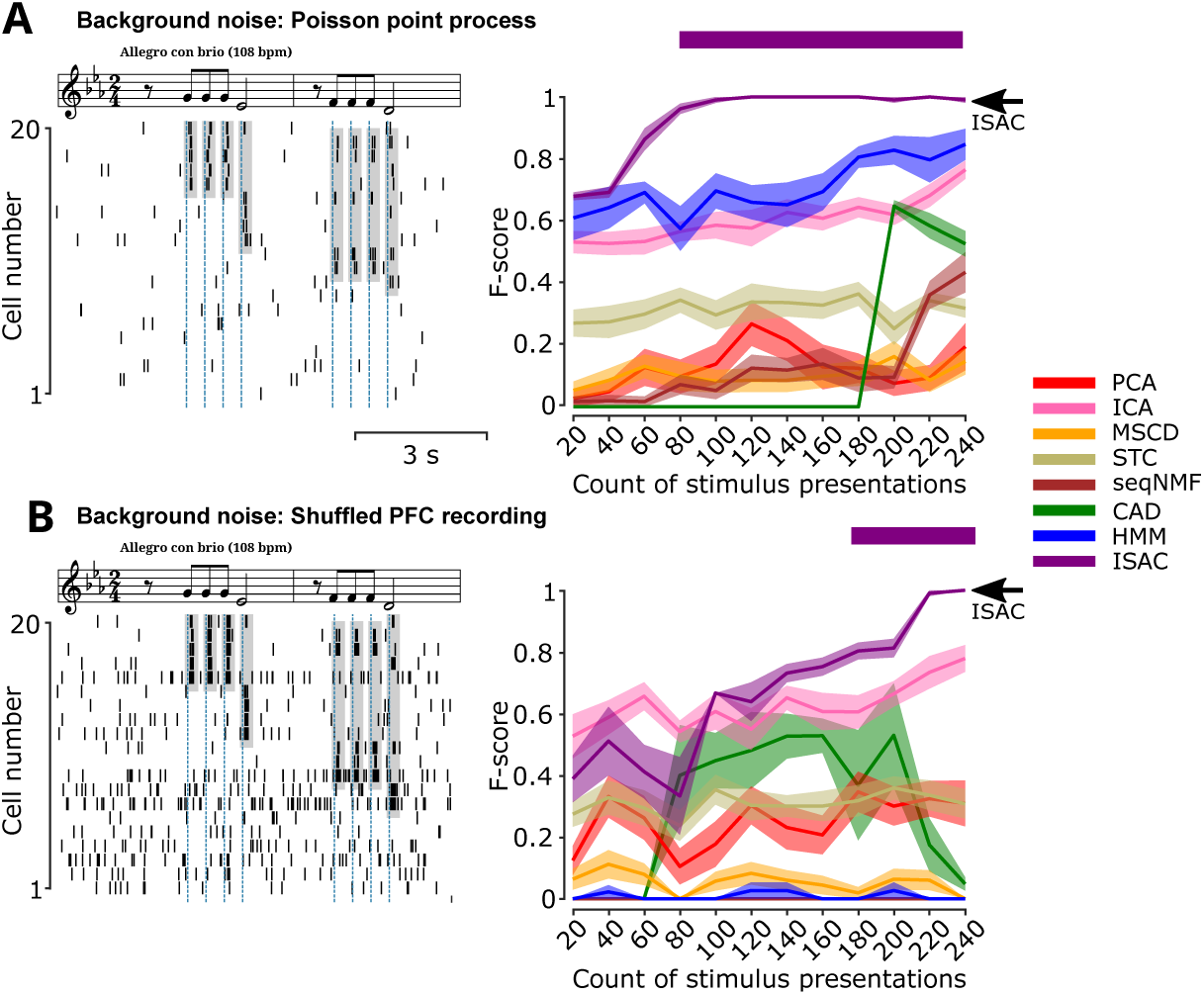
Comparison of respective performances of current algorithms and ISAC on simulated recordings of evoked neuronal responses. (A) Simulated evoked responses of simulated SAC assemblies embedded in Poisson point process background activity. Left: examples of SAC assembly activations to the presentation of the first eight notes of Beethoven’s 5th symphony. Right, same presentation as in Figure 4 but changing the number of auditory events in the dataset (repetition of the eight notes shown). (B) Same as A but using background activity from published experimental recordings of prefrontal cortical neurons with shuffled neuron identities assigned to individual spikes, preserving spiking statistics (see Methods).

As a whole, the poor to moderate performance of most algorithms can be partially accounted for by the fact that they detect partial fragments of SAC assemblies or false positive amalgams of ground truth assemblies. Indeed, when we evaluated the detection of neuron pairs within the ground truth assemblies—ignoring whether the full assemblies were recovered—the algorithms performed notably better, with almost all detected pairs corresponding to true pairs in the ground truth (**Supp. Fig. 7**). These results indicate that, despite their limited success in recovering full simulated SAC assemblies, many of the tested methods correctly identified most pairwise co-activation relationships present in the ground truth.

**Figure 7:**
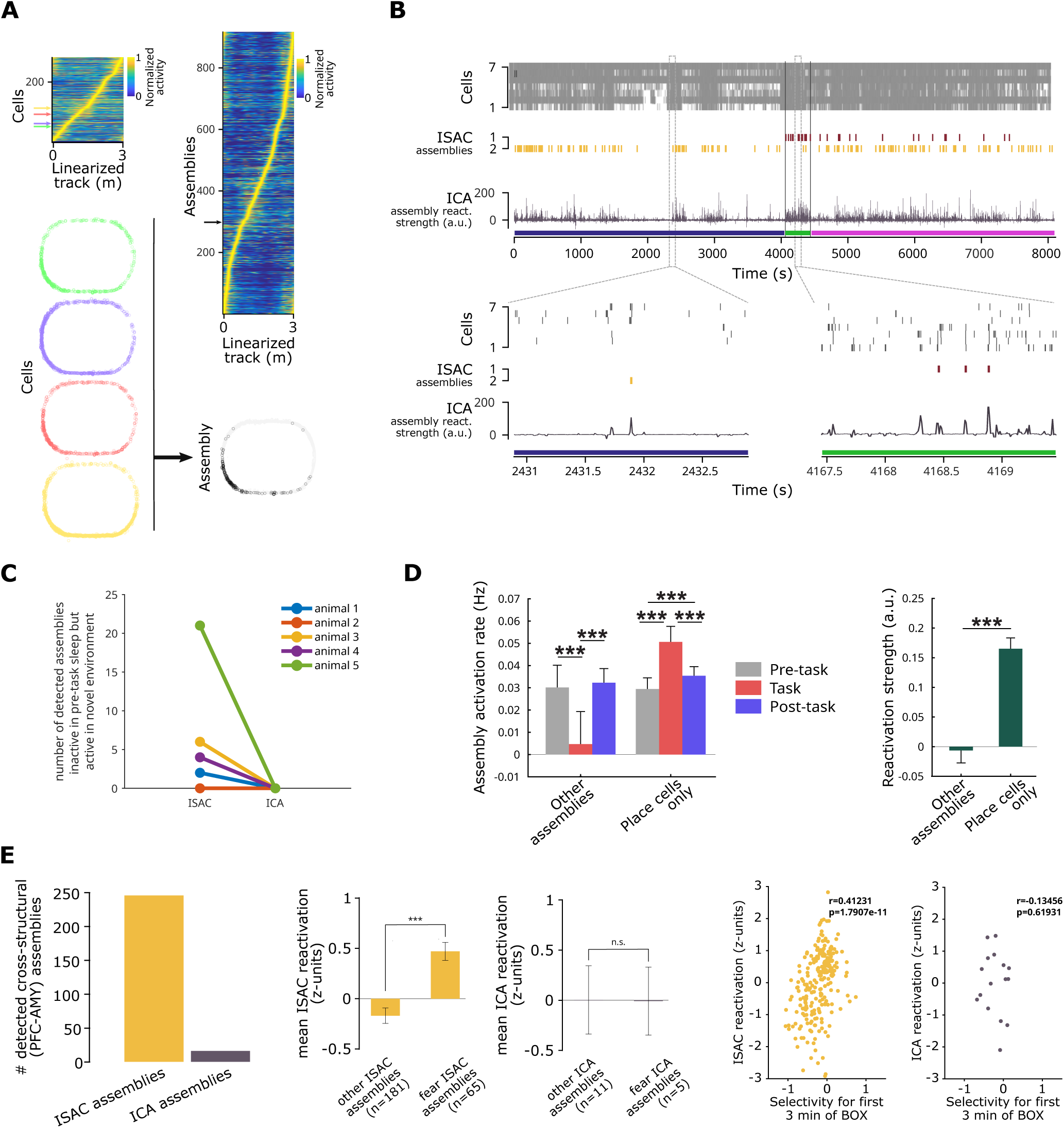
Examples of analytic opportunities unlocked by ISAC. (A) Top Left: all recorded hippocampal cells in a dataset (Drieu et al., 2018) ordered by the spatial position of their firing peak during a task where rats explored a novel environment. Color shows each cell’s firing rate divided by its peak firing rate. Top right: all assemblies identified by ISAC, ordered by the spatial position of their firing peak during the task. Note that assemblies displayed less out-of-field activity, indicating more specific spatial firing. Bottom: example of four cells (denoted by colored arrows in top-left plot) forming an assembly (black arrow in top-right plot) with higher spatial selectivity. (B) Example of assembly activity when the animal is first exposed to a novel context. Colored bars above the time axis indicate the three stages of recording: pre-task rest (blue), task (green), and post-task rest (purple). Top, raster plot of 7 selected cells’ activity during the task. Middle, raster plot showing the activity of two ISAC assemblies. ISAC assembly 1 (red) is silent during pre-task rest, becomes active during exploration, and remains active afterward. ISAC assembly 2 (yellow) is active throughout all stages. Bottom: the ICA component most correlated with ISAC assembly 1 (highest Pearson’s *r*). Panels below show zoomed views of selected time windows. Note that although this ICA component was selected to represent ISAC assembly 1, its activity fluctuations reflect contributions from both ISAC assembly 1 (novel) and ISAC assembly 2 (pre-existing, also correlated with the ICA component), and as a result, the component appears active during pre-task rest despite ISAC assembly 1 being silent. (C) Quantification of the effect illustrated in (B). The number of assemblies silent in pre-task sleep but active in behavior was higher for assemblies detected by ISAC than assemblies detected by ICA (ISAC: 33 such assemblies out of 654 total detected assemblies; ICA: 0 such assemblies out of 55 total detected assemblies; p = 0.0440, chi-square test). (D) Left: Activity of ISAC-detected assemblies during the pre-task, task and post-task periods. Assemblies are divided into two groups: those including non-place cells and those composed exclusively of place cells. Note that only the assemblies composed exclusively of place cells increased their activity in post-task sleep compared to pre-task sleep (\*\*\**p <* 0.001, Wilcoxon rank-sum tests; post-hoc tests with Bonferroni correction). Right: reactivation strength, defined as the relative increase of activity in post-task sleep as compared to pre-task sleep (\*\*\**p <* 0.001, Wilcoxon rank-sum test). (E) Left: Number of prefrontal cortex (PFC) and amygdala (AMY) cross-structural assemblies detected by ISAC and ICA in an example fear recall recording (Pompili et al., 2025). Center: Reactivation strength (increase of post-task rest activity relative to pre-task rest activity) of fear modulated assemblies vs. non fear modulated assemblies with ISAC and ICA. SAC assemblies with high activity in the first 3 minutes of re-exposure to the conditioning context (fear assemblies) showed higher reactivation than other assemblies (\*\*\**p <* 0.001, Wilcoxon rank-sum test). Right: Reactivation strength as a function of firing selectivity for fear context exposure for ISAC-derived and ICA-derived assemblies. Note that the strong relationship between fear-response and reactivation observed for SAC assemblies failed to reach significance in ICA components, which might be due to very low number of ICA components responsive to the fearful environment.

To evaluate performance under more realistic physiological conditions, we used background noise from previously published chronic high-density recordings with shuffled neuron identities assigned to individual spikes. Importantly, this preserves the heterogeneous firing rates and temporal structure, including non-stationary fluctuations, observed in physiological recordings. SAC assemblies were then assigned by randomly selecting cells designated as members, and assembly activations were simulated by selectively shifting specific member spikes in time, creating datasets that were challenging for all algorithms. Although the other algorithms performed poorly across all dataset types, ISAC consistently achieved the best results (**Figure 5**; **Supp. Figs. 8–9**). Importantly, ISAC performance was preserved when assembly activations were restricted to specific periods of the recording, indicating that assemblies can be reliably detected despite non-uniform temporal activation patterns (**Supp. Fig. 10**). Likewise, ISAC did not detect any assemblies in additional control datasets containing no simulated assemblies (n=0), indicating that non-stationary population activity alone does not produce false-positive detections (Supp. Fig. 11).

While the above simulations were tailored to evaluate the performance of algorithms in longer recordings, brief recordings are often employed in research on sensorimotor and decision-making systems and include events that are repeated throughout the recording session. In such cases, where the relevant events are known a priori, restricting analysis to the corresponding epochs may be desirable. To assess assembly detection under these conditions, we simulated an auditory processing experiment with four overlapping simulated SAC assemblies responding to the first eight notes of Beethoven’s 5th symphony. In these event-based simulations, all algorithms’ performance depended on the number of repetitions and therefore on the number of repeated assembly activations. When fewer than 20 stimulus presentations were available, ISAC did not provide an advantage over existing methods, as no algorithm outperformed the others in this challenging regime. As the number of recorded activations increased beyond this range, ISAC consistently outperformed the alternative algorithms in the Poisson point-process simulations (**Figure 6A** and **Supp. Fig. 12A**) and in simulations with realistic spiking statistics (**Figure 6B** and **Supp. Fig. 12B**), demonstrating that its additional statistical constraints become advantageous once sufficient evidence is available. Also in these simulations, the performances were noticeably higher when searching for pairwise synchrony relationships (**Supp. Fig. 13**).

We additionally quantified the computational scaling of ISAC (**Supp. Fig. 14**). Runtime increased approximately linearly with recording duration, while increasing more rapidly with the number of recorded neurons, as expected from the exhaustive search strategy. Nevertheless, ISAC analyzed large datasets (10̃00 neurons) in reasonable time (*<*1 hour), indicating that the algorithm remains practical for offline analyses despite its higher computational cost relative to projection-based methods.

### Application of ISAC to physiological data

The sensitivity and specificity provided by ISAC open doors to new kinds of analyses. Beyond benchmarking detection accuracy, we therefore aimed to illustrate several classes of biological questions that become accessible through improved identification of synchronous cell assemblies. These examples below illustrate several such applications, highlighting how increased sensitivity and specificity facilitate the study of assembly formation, cellular composition, and reactivation dynamics. To use the algorithm with real recorded data, as a case study, we used electrophysiological recordings of hippocampal neurons in rats performing a task in a novel environment (Drieu et al. 2018a). ISAC detected vast numbers of assemblies in this dataset, even surpassing the number of cells in each recording (**Figure 7A**). This effectively expanded the dataset’s representational space, in contrast to conventional dimensionality-reduction approaches. Such expansion and the specificity of the detected assemblies enables the study of assembly formation and dynamics in ways that were previously impossible. For example, a key feature of learning is the formation of novel cell assemblies. Theoretically, such novel assemblies should be silent before task exposure, become active during learning, and persist in their activity afterward. Indeed, we would expect such novel assemblies to emerge during exploration of a new environment. Yet because current techniques often detect multiple SAC assemblies within a single amalgamated group, the activity of one assembly is confounded by the activity of others. As a result, it becomes impossible to identify assemblies that were completely inactive before the task. Consequently, studies of learning are typically limited to tracking assemblies that show only relative increases in activity compared with pre-task sleep, rather than assemblies that are genuinely novel. The specificity of ISAC ensures that distinct groups are not mixed together, and its sensitivity allowed us to identify assemblies that were completely inactive in pre-task sleep, became active during the task, and remained active in post-task sleep (**Figure 7B**). This provides a direct example of a biological analysis enabled by ISAC that is not feasible when multiple assemblies are amalgamated into a single group.

A major difference between ISAC and matrix factorization methods (including the widely used ICA) is that ISAC deterministically defines which cells belong to each assembly. That is, with ISAC, the exact identity of the cells in a group is known. This allows assemblies to be classified based on their cellular composition. For example, in our dataset, we hypothesized that most reactivation during post-task sleep would involve place cells that encoded the novel environment. We therefore compared assemblies composed exclusively of place cells to assemblies that also included other pyramidal cells. Only the exclusively place-cell assemblies showed increased activity in post-task sleep relative to pre-task sleep, whereas the activity of other assemblies remained unchanged (**Figure 7C**). Such analyses rely on the unambiguous assignment of cells to assemblies, a property that is not provided by matrix factorization approaches. Finally, we sought to demonstrate ISAC’s ability to identify cross-structural assemblies (Oberto et al. 2022; Pompili et al. 2026) using previously published simultaneous recordings from the amygdala and prefrontal cortex during a fear-recall test following fear conditioning (Pompili et al. 2025b). Among the 246 ISAC assemblies detected, 65 showed a preferential increase in firing during the first three minutes of re-exposure to the conditioning context (“fear assemblies”), and these assemblies exhibited significantly stronger reactivation during subsequent sleep (**Figure 7E**). This pattern did not emerge when using ICA-derived assemblies, indicating that the larger and more comprehensive set of assemblies detected by ISAC revealed trends that remained inaccessible with ICA’s lower detection precision.

This illustrates how improved assembly detection can reveal biologically meaningful relationships that remain hidden when assemblies are incompletely recovered.

## DISCUSSION

Here we proposed formal criteria for assemblies that ensure they are composed of synchronously active cells, and introduced ISAC, a novel method for the identification of all assemblies fulfilling these criteria within physiological recordings. In simulated datasets where the ground truth is known, ISAC outperforms previous methods by detecting more true positive assemblies while simultaneously detecting fewer false positives. In real-world applications, the ability of ISAC to minimize the detection of erroneously merged groups enables more precise assessment of the functional contributions of cell assemblies. Indeed, it allowed us to pinpoint the formation of assemblies that were active exclusively during and after exposure of the animal to a novel environment. ISAC also makes it straightforward to classify assemblies according to the physiological characteristics of the cells that compose them and to analyze tendencies in their order of firing. The high number of detected assemblies provides statistical power for these analyses, while avoiding cross-pattern mixing and false positive detection.

Unlike commonly utilized methods, ISAC does not rely on a static metric like a pairwise correlation matrix to extract group identities and is thus not constrained in detecting a maximum number of groups. PCA (Peyrache et al. 2010) and ICA (Lopes-dos-Santos et al. 2013) both reduce the dataset into a limited set of factors and, in our tests, detected amalgams of different, unrelated groups within single components. This can obscure unique patterns in the data that may carry selective or informative signals. Avoiding dimensionality reduction entirely allows ISAC to effectively expand rather than compress a dataset by detecting all existing synchronously active cell assemblies. The activity of all these cell assemblies can be directly explored and analyzed – browsed, annotated, and linked to behavior – providing a rich, explicit description of population activity rather than a compressed summary of the data.

A complementary line of research has developed rigorous statistical frameworks for quantifying higher-order interactions in neuronal populations (Amari 2001; Shimazaki et al. 2012; Stella et al. 2019; Torre et al. 2013). These approaches are closely related to ISAC’s unity criterion, which tests whether the synchronous activity of a group of neurons can be explained by the activity of its subsets, and therefore shares an important conceptual connection with the study of higher-order correlations (HOC). However, while HOC frameworks focus on quantifying higher-order interactions, ISAC is designed to specifically identify the cellular composition of synchronously active assemblies that satisfy explicit affinity and unity criteria.

As an exhaustive search algorithm, ISAC is computationally more demanding than projection-based methods such as PCA or ICA. This reflects a deliberate trade-off between computational efficiency and exhaustive identification of synchronously active groups. As computational resources continue to improve, we believe this trade-off is increasingly acceptable for offline analyses where accurate assembly identification is the primary objective.

Other methods to detect putative cell assembly proxies have overcome many limitations of matrix factorization methods, but have their own constraints. In the case of CAD (Russo and Durstewitz 2017), membership of only the most recently added cell within a pattern is validated. The lack of cross-validation against all members within that pattern can introduce false positives into detected assemblies, as we have shown (Fig. 1c, Supp. Fig. 1A). Strong correlations of a cell with part of a pattern can therefore lead to erroneous detection. HMM (Maboudi et al. 2018) exhibits high accuracy when the exact number of cell assemblies is provided by the user; however, small deviations from the optimum value lead to rapid deterioration of performance. This is a major hurdle for recorded data, as the ground truth is unknown and estimating the number of assemblies within a dataset is difficult.

The shuffled data-derived simulations in Figure 5 were accompanied by a dramatic drop in the performance of some algorithms compared with the stationary Poisson process simulations in Figure 4. The reduced performance of PCA and ICA is consistent with the departure from the assumption that assembly-related covariance dominates the correlation matrix. In recordings with heterogeneous, non-stationary population activity, global fluctuations contribute substantial covariance unrelated to cell assemblies, reducing the fraction of variance explained by the injected assemblies. Similarly, the HMM implementation used here first identifies population burst events and searches for assemblies only within those events. Because our simulation consisted of a background of realistic fluctuations in population firing with assembly activations injected at random times, some ground-truth assembly activations occurred outside detected global population bursts, reducing HMM sensitivity under these conditions. By contrast, ISAC evaluates each neuron’s coupling to a candidate assembly relative to the surrounding neuronal population, making it less sensitive to global rate fluctuations and non-stationarities.

It is important to note that, although ISAC outperformed the other methods in detecting synchronously active cell groups, most methods correctly identified the pairwise co-activation relationships present in the ground truth (Supp. Fig. 7; Supp. Fig. 9). This suggests that the performance differences may reflect distinct objectives of the methods rather than their general ability to detect neuronal population structure. In particular, methods designed to identify different forms of population activity may perform equally well or better when assemblies are defined according to alternative objectives rather than synchronously active groups. Our results therefore demonstrate ISAC’s effectiveness in recovering synchronously active cell groups as defined by the affinity and unity criteria, rather than its general superiority for analysing neuronal population activity.

Our benchmark necessarily represents a selection rather than an exhaustive comparison of all existing assembly-detection approaches. We selected methods that had been independently implemented beyond their originating groups, with the aim of focusing on approaches adopted by the broader community. This criterion excludes some potentially relevant methods (e.g. Muiño and Borgelt 2014), and our comparisons should therefore not be interpreted as a comprehensive evaluation of all available methods.

Overall, ISAC addresses the limitations of existing assembly-detection approaches by explicitly searching for groups of synchronously active cells (SAC assemblies) by evaluating them against the affinity and unity criteria. These two criteria ensure the detection of cells that are significantly active together as a collective unit, avoiding the mixing of groups within a single pattern. To the best of our knowledge, ISAC is the only method which specifically seeks to disentangle the identity of individual groups that may participate in larger assemblies. This sets the ground for further work on the functional role of cell assemblies in cognitive processes, on dynamic interactions within and between cerebral subsystems, and the fundamental aspects of brain computation.

## METHODS

### Criteria for synchrony

The identification of groups of synchronously active cells (SAC assemblies) relies on two complementary statistical criteria: *affinity* and *unity*. These criteria are evaluated from spike timing data within a user-defined window of synchrony (Δ*t*, set to 30 ms unless otherwise noted).

Importantly, spike trains are not discretized into temporal bins. Instead, synchrony is evaluated directly from the continuous spike times: spikes from different neurons are considered part of the same activation event when they occur within Δ*t*. This avoids artefacts associated with assigning spikes to fixed, non-overlapping temporal bins.

The *affinity* criterion assesses whether each candidate neuron in a group preferentially fires with the group’s activations compared to how the rest of the neuronal population activity (non-member population activity) fires with the group’s activations. For the candidate group *X* to pass the affinity criterion, all candidate member neurons must individually meet this condition, ensuring that each member neuron within the group increases its firing rate when the group is active beyond what can be expected from fluctuations in the global population firing rate. In particular, if neuron *A* is part of group *X*, then *A* preferentially fires with the other neurons of group *X*, and this preference goes beyond what could be expected from global fluctuation statistics impacting the whole population.

To test this for a given neuron *A*, activations are computed as moments when all other members of *X* (disregarding *A*) are active together within the window of synchrony Δ*t* (that is, time windows in which all other members of *X* emit at least one spike each). The number of spikes *a*_sync_ fired by *A* within Δ*t* of all these activations is computed. To estimate global population fluctuations, an equivalent value is computed for the non-member population activity (defined here as the pooled spike train formed by all neurons not belonging to the candidate group), yielding *Pop*_sync_. The corresponding proportions are

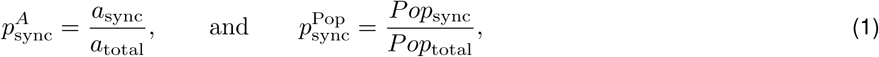

where *a*_total_ and *Pop*_total_ are the total number of recorded spikes by *A* and by the non-member population activity, respectively. The affinity criterion for neuron *A* is satisfied if according to a two-proportion *z*-test.

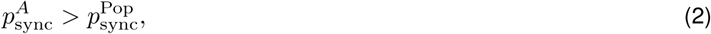

The *unity* criterion assesses whether each candidate neuron in a group contributes disproportionately when all other members of the group are active together, compared to when only a subset of the other members is active. To test this for a given neuron *A*, let *m* be the number of other neurons in *X*. Activations of the entire group are computed as moments when all *m* other members are simultaneously active within the window of synchrony Δ*t*. The number of spikes *a*_sync_ fired by *A* within Δ*t* of these full-group activations is computed (same as above). Then, for each subset of *X* containing *m −* 1 members, the number of spikes fired by *A* within Δ*t* of the subset’s activations is computed. Because different subsets may have different total time covered by their Δ*t* windows (due to differing activation counts and possible overlaps between windows), we normalize each subset’s spike count by multiplying it by the ratio of the full-group activation duration to the subset activation duration. This yields the normalized count *a*_sub_, which can be directly compared to *a*_sync_. The corresponding proportions are

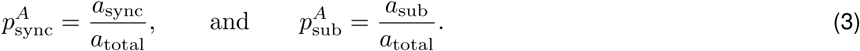

The unity criterion for neuron *A* is satisfied if

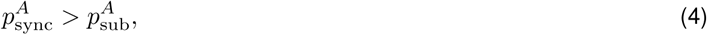

according to a two-proportion *z*-test for all subsets of size *m −* 1. For the candidate group *X* to pass the unity criterion, all candidate member neurons must individually meet this condition, ensuring that the group functions as a cohesive unit rather than as loosely correlated subsets.

For both criteria, the two-proportion *z*-tests therefore compare

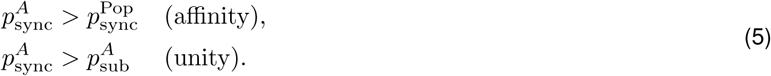

The corresponding *z*-values are considered significant when they exceed the user-defined threshold *Z*. In this manuscript, *Z* is set to 2.57, equivalent to *p <* 0.01, unless otherwise noted, corresponding to an estimated false-positive rate of 0.2% (**Supp. Fig. 4**).

### Iterative detection of SAC assemblies

We developed ISAC (Identification of Synchronously Active Cells) to find all SAC assemblies (Synchronously Active Cell assemblies) that satisfy the affinity and unity criteria. The procedure starts with every possible combination of three neurons in the recording. Each triplet is tested for synchrony using the affinity and unity criteria as described above. A triplet that satisfies both criteria is defined as a SAC assembly. Then, for each significant three-cell SAC assembly, ISAC attempts to expand it by testing all possible four-cell groups that would be formed by adding one additional neuron to the existing triplet. Each four-cell group is then assessed for affinity and unity. For instance, if a triplet ABC satisfies the criteria, all 4-cell combinations containing ABC, such as ABCD, ABCE, ABCF (etc.…), are evaluated against the affinity and unity criteria in the subsequent cycle. If both criteria are satisfied, the newly expanded group is defined as a SAC assembly. This process continues iteratively, expanding n-cell SAC assemblies to n+1 cell groups, until no further significant expansions can be made. Additionally, ISAC can be optionally configured to identify SAC assemblies that are distributed across multiple brain regions (Figure 7E). The user can optionally provide a mapping of each neuron to its anatomical region and set a minimum number of distinct regions required for a SAC assembly. When a candidate n-cell SAC assembly has not yet met the minimum region criterion, the search space for the n+1 expansion is restricted exclusively to neurons from brain regions not yet represented in the candidate group. Once the minimum number of regions is satisfied for a given SAC assembly, this constraint is lifted, and the algorithm proceeds to test the addition of any statistically relevant neuron, regardless of its region, as described previously.

When either affinity or unity fails to reach significance, the expansion process halts. An optional parameter, the relevance threshold R, allows the user to filter SAC assemblies and detect only the ones that are active at least R times in the dataset (in this manuscript, R is set to 1).

### Merging of SAC assemblies

Following iterative detection, ISAC includes an optional defragmentation step designed to recover large assemblies that may have been identified as separate fragments. This process merges detected n-cell fragments (for example, ABC, ABD, ACD, and BCD) into larger n+1-member assemblies (for example, ABCD) when the resultant larger assembly satisfies the affinity criterion. A user-defined tolerance parameter, D, sets the maximum allowable percentage of missing fragments (that is, the fraction of the (*_n_^n^*^+1^) possible *n*-cell sub-assemblies of the merged assembly that may be missing from the set of detected SAC assemblies). ISAC performances relative to tolerance values are shown in **Supplementary Figs. 15-16**. In the analyses presented in this manuscript, D was set to 10%.

### ISAC code availability

The code to run ISAC can be found in the following GitHub repository: https://github.com/mnpompili/ISAC.

### Selection of Currently Used Methods for Cell Assembly Identification

To form an inventory of existing methods for cell assembly detection in electrophysiological recordings, we conducted a literature search that returned 26 research articles describing algorithms for the detection of multi-cellular patterns within spike trains. We then focused on methods that were successfully implemented in experimental works by at least one lab different from the one that developed the algorithm. This criterion was chosen to focus the benchmark on methods that had been adopted by the broader research community and demonstrated use beyond their originating groups, without selecting according to any particular methodological family. Seven methods were identified as meeting these selection criteria: Principal Component Analysis (PCA)(Peyrache et al., 2010), Spike-Train Communities (STC) (Humphries, 2011), Independent Component Analysis (ICA) (Lopes-dos-Santos, Ribeiro and Tort, 2013), Markov Stability for Community Detection (MSCD) (Billeh et al., 2014), Cell Assembly Detection (CAD) (Russo and Durstewitz, 2017), Hidden Markov Model (HMM) (Maboudi et al., 2018), and Sequential Non-negative Matrix Factorization (seqNMF) (Mackevicius et al., 2019). We excluded the Louvain method (Blondel et al. 2008), a generic clustering algorithm constituting the backbone of MSCD as well as another method based on hidden Markov models (Mazzucato et al. 2015), due to its similarity to the HMM method described by Maboudi et al. (2018).

Many methods return cell assemblies as a weighted vector of components, with all recorded units participating to some extent in each component (PCA, ICA, HMM, seqNMF). To identify cell assembly members for these methods, we set a threshold on the units’ participation weights, as previously described (Oberto et al. 2022; Trouche et al. 2019; Van de Ven et al. 2016). Component weights were z-scored and cells whose z-scored weight surpassed 2 were considered members of the detected group.

### Implementation of Currently Used Methods

A complete list of the parameters used for each algorithm, together with their source and any ground-truth information provided during the benchmark, is provided in **Supplementary Table 2**. **Principal Component Analysis (PCA) (Peyrache et al., 2010)** was implemented using MATLAB code in the FMAToolbox (http://fmatoolbox.sourceforge.net). Briefly, the dataset comprising N neurons was binned into T millisecond windows, with two successive bins overlapping by T/2 milliseconds. The binned spike matrix was Z-scored and a pairwise Pearson’s correlation matrix C was constructed. We ran eigendecomposition on the obtained matrix C and eigenvectors whose eigenvalues surpassed the upper threshold of the Marcenko-Pastur distribution were retained as significant components. For runs on simulated datasets, T was set to 30 milliseconds, corresponding to the maximum temporal jitter applied to spikes participating in an assembly activation and was equal to the ISAC window.

**Spike-Train Communities (STC) (Humphries, 2011)** was carried out using MATLAB code from the official GitHub repository: (https://github.com/mdhumphries/SpikeTrainCommunitiesToolBox). The quantization values of the convolution function were set to 1 millisecond, the standard deviation of the Gaussian convolution function was set to 15 milliseconds, and pairwise correlations were computed using the cosine function.

**Independent Component Analysis (ICA) (Lopes-dos-Santos, Ribeiro and Tort, 2013)** was implemented using MATLAB code in the FMAToolbox (http://fmatoolbox.sourceforge.net). Briefly, we z-scored the spike matrix M and projected it onto its K significant principal components, detected as described for the PCA method above. We then ran ICA using the FastICA package for MATLAB (http://research.ics.aalto.fi/ica/fastica) on the projected matrix to obtain weighted independent components.

**Markov Stability for Community Detection (MSCD) (Billeh et al., 2014)** was implemented using MATLAB code from the official GitHub repository (https://github.com/CellAssembly/Detection). The code was optimized by using a spike template instead of discretizing an exponential decay function for each spike. This optimization yielded results identical to the original code but with reduced temporal complexity. The binless window step was set to 1 millisecond, and the time constant of the exponential profile was set to 3 milliseconds. Markov time consisted of 81 values following an exponential scale (base 10) between 0.01 and 100. The optimal Markov time was determined as described by Billeh et al. (2014), that is as the longest plateau in the number of communities when varying Markov time.

**Cell Assembly Detection (CAD) (Russo and Durstewitz, 2017)** was implemented using MATLAB code from the GitHub repository (https://github.com/DurstewitzLab/Cell-Assembly-Detection). For all runs on simulated datasets, the bin size was set to 30 milliseconds, and the maximum lag variable was set to 2. The significance level (alpha) was kept at the default value of 0.05.

**Hidden Markov Model (HMM) (Kourosh Maboudi et al., 2018)** was implemented using Python code from the official GitHub repository (https://github.com/nelpy). Data was binned into 30 millisecond windows. Population burst events were used to detect components and were detected as described by Maboudi et al., 2018 as bins where multiunit activity surpassed three standard deviations. As the algorithm does not estimate the number of components, we used three different approaches for setting the number of assemblies in the simulations. For one-third of the simulations, we used the ground truth number of cell assemblies. For another third, we set the number at 50% of the ground truth value. In the final third, we used twice the ground truth number of assemblies.

**Sequential Non-Negative Matrix Factorization (seqNMF) (Mackevicius et al., 2019)** was carried out using MATLAB code from the GitHub repository (https://github.com/FeeLab/seqNMF). Data was binned into 30 millisecond windows. The number of factors K was set to the number of SAC assemblies in the ground truth dataset. Where specified, optimal K was determined by minimizing the median dissimilarity measure, computed as described in Mackevicius et al. (2019), and ran on values of K between 1 and 10. Since there was no preference to cell order in SAC assemblies, the length of each factor was set to one.

### Detection of putative cell assembly activations

The all-or-none activations method identifies discrete, binary events corresponding to the activations of putative cell assemblies. This approach is applicable when cell assemblies have been defined using binary membership vectors that specify whether each neuron is a member (1) or not (0) of a given assembly. An assembly activation is defined as any moment in time (within a time window Δ*t*) during which each member fires at least one spike. By requiring the simultaneous spiking of all assembly members, this method provides a stringent criterion for activation, ensuring high specificity. However, it may overlook instances where only a subset of the assembly is active, potentially missing partial or subthreshold activations. Importantly, this approach does not require spike trains to be discretized into temporal bins. Instead, synchrony is evaluated directly from the continuous spike times: spikes from different neurons are considered part of the same activation event when they occur within Δ*t*. This avoids artefacts associated with assigning spikes to fixed temporal bins.

An alternative to the above is the reactivation strength method, which quantifies the degree to which a putative cell assembly is active at a certain time bin (used for ICA in Figure 7). We compute reactivation strength as described in Peyrache et al. (2010). The reference assembly (e.g., component) is represented as a weighted vector where weights correspond to each neuron’s contribution to the assembly. To emphasize co-activity among neurons and suppress the influence of individual firing rates, the weighted vector is squared, and the diagonal of the resulting matrix is set to zero – effectively nullifying self-contributions. Spike trains are binned, and the binned spike matrix is z-scored across time for each neuron. The reactivation strength at each time bin is then calculated by projecting the z-scored spike data onto the modified weighted vector. This projection yields a reactivation strength value for each time bin. The resulting time series of reactivation strengths is subsequently z-scored. Significant epochs are defined as moments when the z-scored reactivation strength exceeds a threshold of 5. Reactivation strength was used for the detection of the activation of components detected by ICA in Figure 7.

### Construction of simulated datasets

The simulation framework followed established benchmarking approaches for neuronal assembly detection algorithms by generating background neuronal activity and randomly embedding activations the activation of ground-truth assemblies throughout the dataset. Assembly activations were simulated by injecting jittered synchronous activity across multiple neurons (Billeh et al. 2014) into an artificial background while compensating for each injected spike by removing a background spike, thereby preserving each neuron’s mean firing rate (Russo and Durstewitz 2017).

We included two different forms of background activity. For the simulations shown in Figure 4, background neuronal activity was generated as a stationary Poisson process, as in previous studies (Lopes-dos-Santos et al. 2013; Russo and Durstewitz 2017; Stella et al. 2019) with an identical mean firing rate of *λ* = 2 Hz for all neurons. Because real-world recordings violate many of the assumptions underlying this idealized background model, including the homogeneous firing rates and the stationarity of the population firing rate, we also performed simulations under conditions that more closely resembled experimental data. Specifically, for the simulations shown in Figure 5, background activity was derived from an experimentally recorded prefrontal cortex dataset ((Peyrache et al. 2018); session 181020) by randomly permuting (i.e., shuffling) neuron identity labels across recorded spikes. This procedure preserved each neuron’s heterogeneous firing rate distribution as well as the non-stationary population-rate fluctuations characteristic of experimental recordings while disrupting the original pairwise correlations between neurons. Consequently, these simulations provided a substantially more challenging and physiologically realistic benchmark for evaluating assembly detection performance.

Between simulations, we varied: the number of SAC assemblies in the dataset (*SAC_number_*) and the number of cells within a SAC assembly (*SAC_size_*), and the SAC assemblies activation rate (*SAC_activations_*). Activations times were randomly distributed throughout the dataset. During each SAC assembly activation, neurons assigned as SAC assembly members fired a number of spikes proportional to their respective firing rate (e.g. 2 spikes for a neuron with a firing rate of 2 Hz, with a minimum of 1 spike). Spikes were jittered within a 30 ms window around the activation time to simulate natural variability, consistent with the gamma rhythm (Buzśaki, 2010). To ensure that the addition of assembly activations did not overly inflate a neuron’s firing rate, for each spike that was added during an assembly activation, a spike was randomly removed from the background activity of the same neuron.

In the baseline configuration, a simulated recording of 50 cells included *SAC_number_*= 20 randomly assigned SAC assemblies, each containing *SAC_size_* = 5 cells. Note that individual neurons could participate in multiple SAC assemblies and each SAC assembly was unique and not a subset of another. The simulation ran for a total duration of *L* = 60 minutes, with *SAC_activations_*= 2 per minute occurring at random timestamps in the dataset. To assess the scalability and adaptability of the methods, *SAC_number_*(10 to 200 assemblies), *SAC_size_*(3 to 15 cells), *SAC_activations_*(0.05 to 10 activations per minute), and dataset length *L* (10 to 100 minutes) were systematically varied in separate simulations. For each simulation, other parameters were held constant to isolate the effects of the tested parameter. Details of all simulations are provided in Supplementary Table 1. Each condition was tested on 15 separate datasets, except for the CAD method, which was tested only twice per condition due to computational demands.

### Statistical evaluation of methods performances

Assembly timescale of the simulated ground-truth assemblies was provided to all methods. All other parameters were taken from the original publications for each method or their public implementations (see **Supplementary Table 2**). To evaluate the performance of the cell assembly detection methods, we used the F-score (Chinchor 1992), which takes into account the accuracy of detected assemblies (precision) and the completeness of detection (recall). The F score is calculated as:

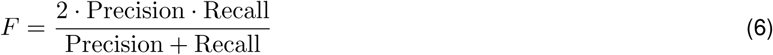

Precision measures the proportion of correctly identified SAC assemblies out of all groups detected by a method and is calculated as:

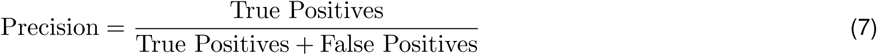

Recall measures the proportion of ground-truth SAC assemblies correctly identified by the method and is calculated as:

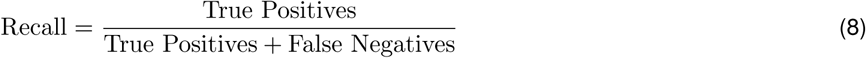

True positives are groups detected by an algorithm that exactly match a ground-truth SAC assembly. An exact match means that all neurons in the detected group correspond to those in a ground-truth SAC assembly, with no extra or missing neurons. False positives are detected groups that do not correspond to any ground-truth SAC assembly, or which are not part of any SAC assembly. False negatives are ground-truth SAC assemblies that were not detected by the method.

Statistical comparisons of F-scores across methods were performed using Wilcoxon’s signed-rank test. In addition to the categories defined above, we quantified sub-categories of false positives based on their relationship with ground truth assemblies for the analyses in Supp. Figures 6, 8, and 12. “Partial ground truth assemblies” were defined as detected groups that include only a subset of neurons from a ground-truth assembly. For example, if a ground-truth assembly consists of 5 neurons and the detected assembly includes only 3 of these neurons (and no other members), it is considered a partial detection. “Mixed ground truth assemblies” were defined as groups that include neurons from at least two different ground-truth assemblies. This occurs when the method incorrectly groups neurons from separate ground truth assemblies into one detected assembly. Finally, “Non-synchronous groups” were defined as groups that do not correspond to any ground-truth assembly and are not mixtures of multiple ground-truth assemblies.

To assess the respective methods’ ability to detect co-firing relationships between pairs of neurons within assemblies, we conducted a pair-level analysis in Supp. Figures 7, 9, and 13. For each ground-truth SAC assembly and each detected group, we extracted all possible unique pairs of neurons. True positive pairs are pairs found in both the detected groups and the ground-truth SAC assemblies. False positive pairs are pairs found in the detected groups but not present in any ground-truth SAC assembly.

### Experimental procedures

Some ISAC’s potential applications were showcased on on-line archives of data from electrophysiological recording in the rat hippocampus (Drieu et al. 2018a) and of the amygdala and prefrontal cortex (Pompili et al. 2025a).

For the hippocampal recordings, briefly, they were on average three to four hours long and consisted of 90 minutes of sleep (“pre-task rest”), followed by a ten-minute task, and then 90 minutes of post-task sleep (“post-task rest”). The task consisted of the animal walking on a treadmill while being transported on a train traveling around a stadium-shaped rail (see original publication for details on experimental procedures). During the task, theta-sequences formed, making this task particularly suited to study the organization and ordering of cell activity within assemblies (Drieu et al. 2018b). Furthermore, the large number of laps conducted by the animal allows for the analysis of place-related activity, while sleep recordings allow us to assess the cell assembly activity after exposure to this environment. Data was limited to the first day of exposure to the environment, enabling us to study the creation of novel task-related assemblies during the animal’s first encounter with the environment. This analysis was applied separately to data collected from 5 rats.

SAC assemblies were extracted using ISAC with Δ*t* = 150 *ms*, reflecting the theta scale observed during active wakefulness. Activations of SAC assemblies detected by ISAC were defined as moments in time where all cells composing an assembly co-fired within a time window of 150 ms (all-or-none activations, see above). Activations of putative cell assemblies detected by ICA were detected using the reactivation strength method (see above). Linearized place fields were created using the function FiringCurve from the FMAToolbox.

For the prefrontal cortex-amygdala recordings, briefly, simultaneous high-density electrophysiological recordings were obtained from the medial prefrontal cortex and amygdala of four rats across five recording sessions per animal (Pompili et al. 2025b). Recording sessions comprised extended sleep periods interleaved with behavioral sessions involving fear conditioning and extinction, with sleep epochs detected using cortical spindle activity to discriminate them from freezing behavior as previously described (Pompili and Todorova 2022). The simultaneous recording of large neuronal populations from these reciprocally connected structures makes this dataset particularly suited to investigate cross-structural cell assemblies, allowing us to identify assemblies containing both prefrontal and amygdalar neurons and examine their activity across behavioral and sleep periods.

For all experimental datasets, including the dataset used in Figure 2 and to generate the complex background for the Figure 5 simulations (Peyrache et al. 2018), we analyzed the publicly available spike-sorted data provided by the original authors. No additional preprocessing of the spike trains (e.g., trial selection, temporal binning, or trimming of the recordings) was performed.

## ACKNOWLEDGMENTS

We thank Michaël Zugaro for discussions on the analyses and comments on the manuscript; Céline Drieu and Adrien Peyrache for making available the hippocampus and prefrontal cortex recordings, respectively; Daniel Bennequin for discussions on an earlier version of the algorithm; and Cédric Colas and Noé Hamou for useful comments on the code. This work was funded by a FFRE grants to MNP and ANR grant to RT. GM is funded by Marseille Neuroschool.

## AUTHOR CONTRIBUTIONS

Project conception and management: MNP, RT, GM. Assembly detection design: RT, GM, MNP. Assembly detection code implementation: RT, GM. Simulated datasets and testing of methods design: GM, RT, MNP. Simulations implementation: GM. Implementation and testing of other methods: GM. Design of analyses of real recordings with ISAC: RT, MNP. Implementation of these analyses: RT, GM. Visualization and writing: MNP, GM, RT, SIW.

---

**Supplementary Figure 1:**
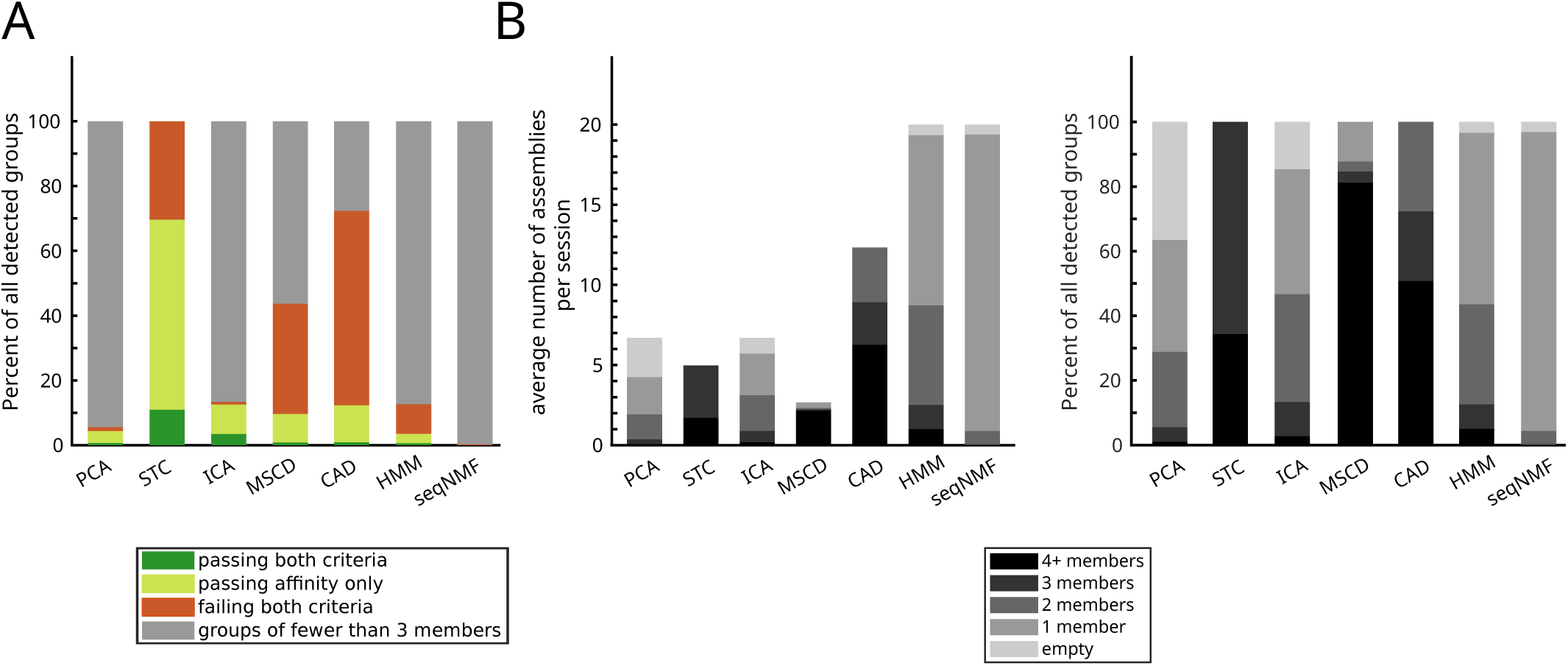
Characteristics of putative cell assemblies detected by different algorithms on recordings of the prefrontal cortex (same dataset of. Fig. 2).. (A) Percent of detected putative cell assemblies passing the affinity and unity criteria. (B) Counts of detected putative cell assemblies of different sizes detected by the different methods. Left: Average number of detected putative assemblies. Right: same data as left, shown as a percentage of all detected assemblies by a given algorithm.

**Supplementary Figure 2:**
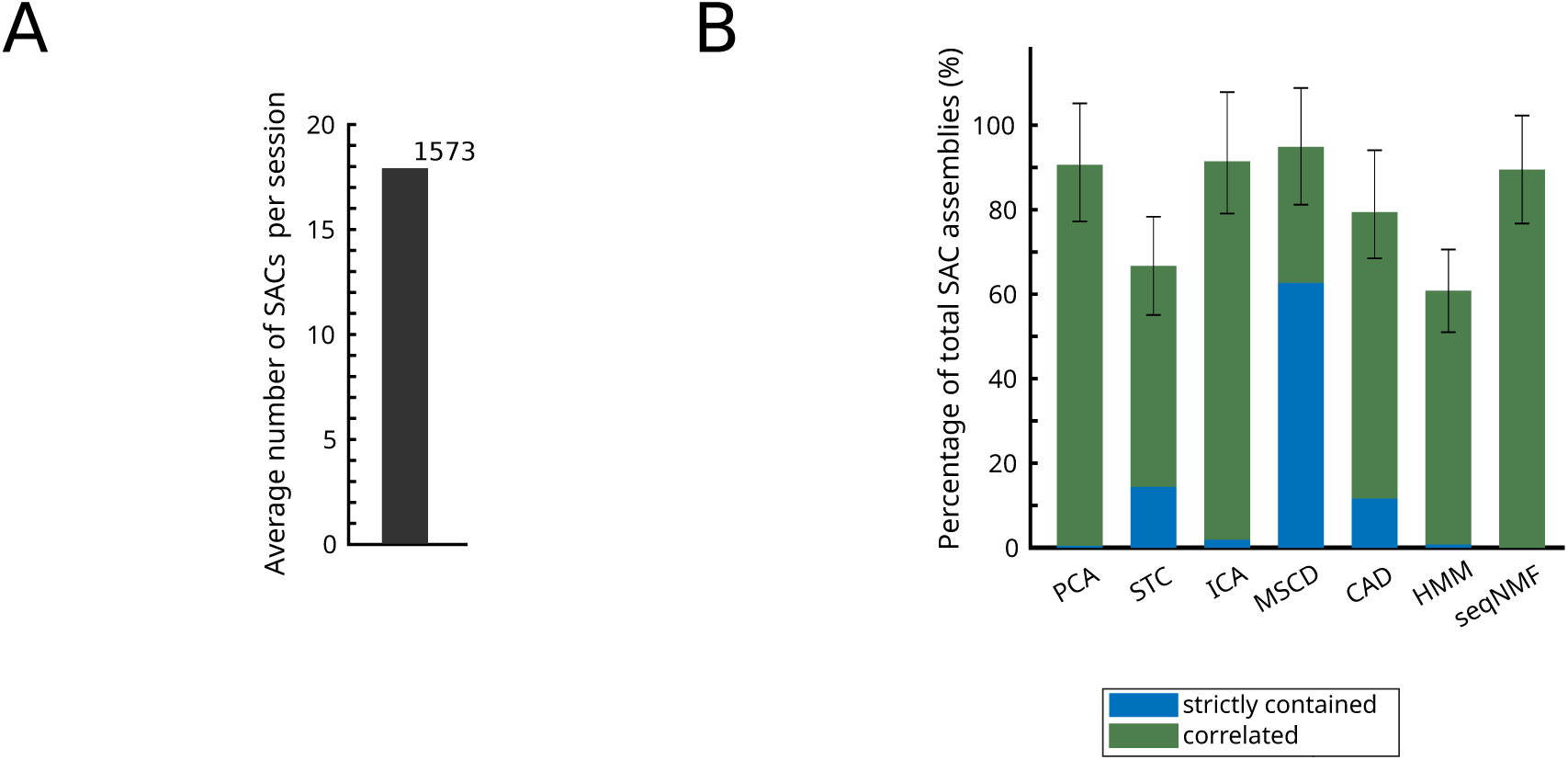
SAC assemblies of 3 members on the prefrontal cortex dataset shown in. Fig. 2 **and Supp. Fig. 1.** (A) Triplets fulfilling both synchrony criteria identified in the prefrontal dataset used in Fig. 2. Bar: average number of SAC triplets per session. The total number of SAC triplets identified for all sessions is indicated above the bar. (B) Relation of the identified SAC triplets with putative assemblies detected by current methods. On average, around 80% of SAC triplets were either significantly correlated (Pearson correlation *p <* 0.05) or contained within putative cell assemblies detected by other methods.

**Supplementary Figure 3:**
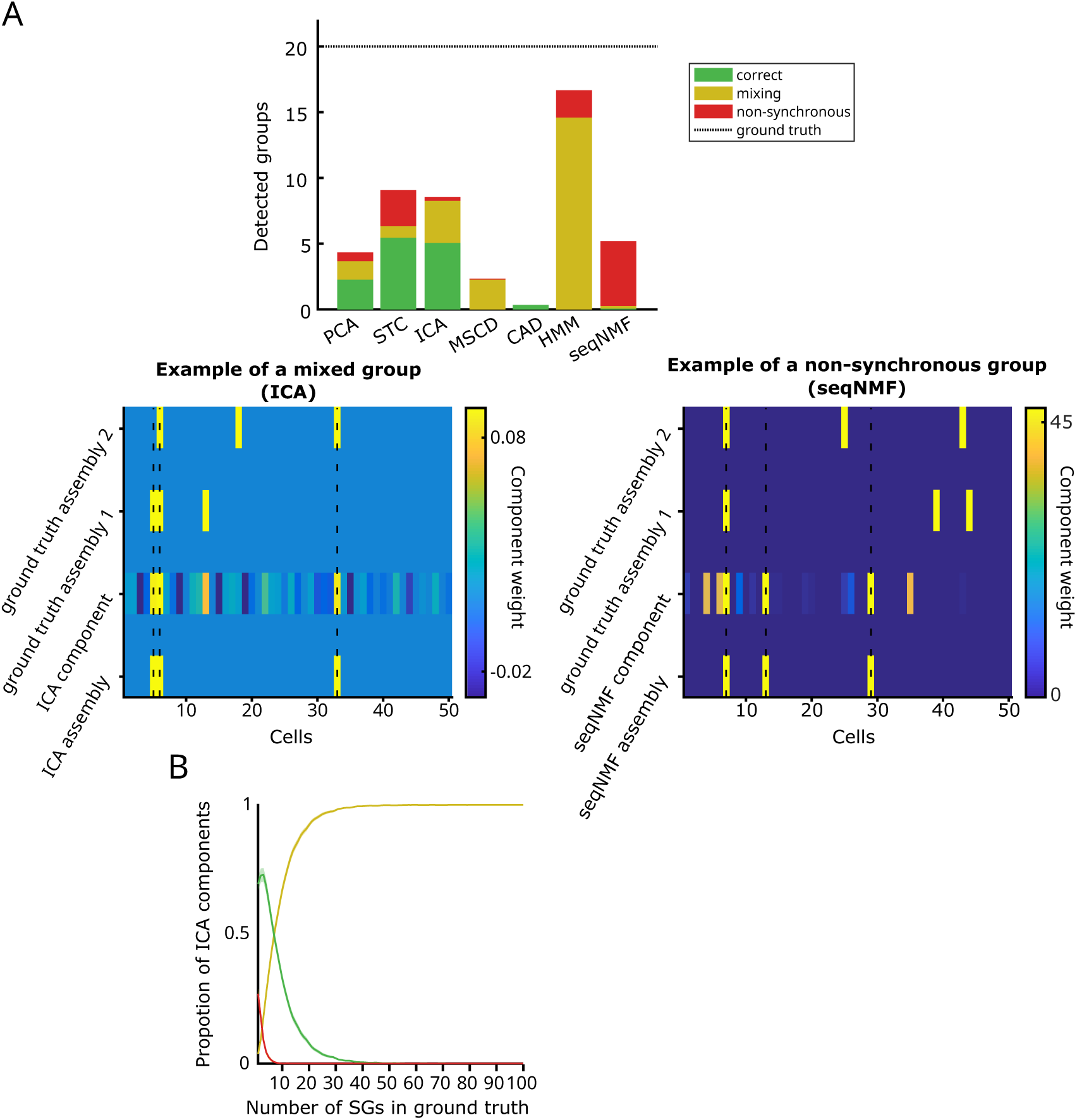
Cell assembly detection on a simulated dataset with twenty SAC assemblies composed of three cells each. (A) Performance of the different methods evaluated as exact groups, mixed groups, and non-synchronous groups. Exact groups are defined as exact matches of groups in the ground truth. Mixed groups are defined as groups whose identity could be explained by pairwise correlations within ground truth groups (between pairs of members). Non-synchronous groups were defined as groups where the inclusion of at least one member cell could not be explained by pairwise correlations within the ground truth. Note that patterns with two or fewer cells were discarded. Top: Average number of detected assemblies. Bottom left: An example of a mixed group detected by ICA. Note how the mixed group is comprised of member cells 5, 6, 13, and 33. While an exact match of this group is not present in the ground truth, it mixes pairs of cells present within two other ground truth assemblies. Cells 6 and 33 are present within ground truth assembly 1, and cells 5, 6 and 13 within ground truth assembly 2. Bottom right: An example of a non-synchronous group detected by seqNMF. The ground truth assemblies with most overlap with the seqNMF members are shown. In the non-synchronous group, the pairs [7, 13], [13, 29], and [7, 29] are not included within any ground truth group. (B) Detection of ground truth assemblies within ICA components, plotted as a function of the number of ground truth assemblies in the ground truth dataset. As the number of ground truth assemblies increases, the proportion of mixed groups also increases, while the proportion of correct groups decreases. For datasets with more than fifty ground truth assemblies, no components represented only one ground truth assembly from the underlying dataset. Color same as in A.

**Supplementary Figure 4:**
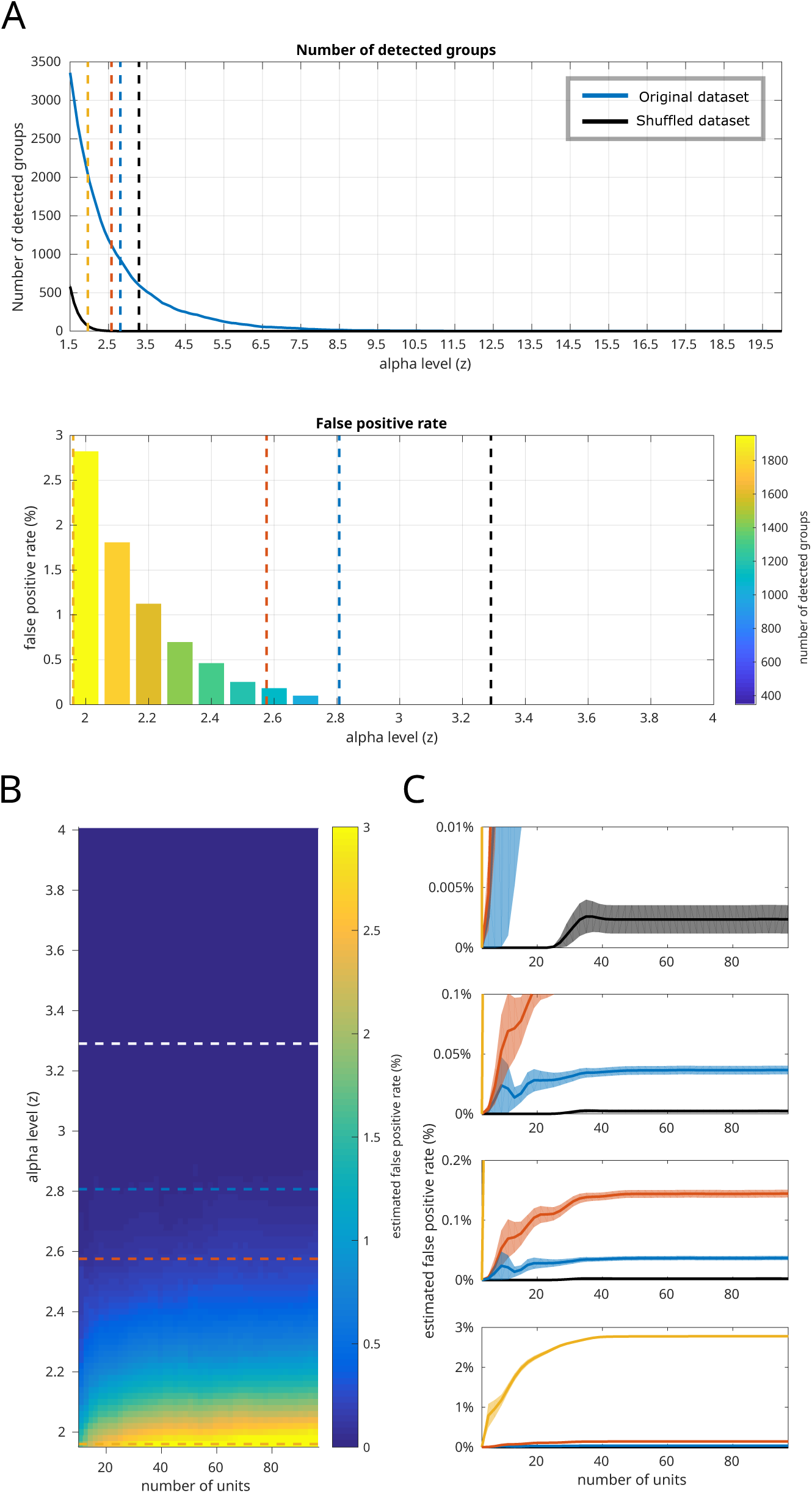
False positive groups detected by ISAC on a dataset recorded in the prefrontal cortex as a function of alpha level. **Z.**. (A) The number of detected SAC assemblies as a function of the alpha level used in ISAC (expressed as a standard normal deviate, the Z-statistic). Top, Blue line: Number of SAC assemblies detected in the original dataset. Black line: Number of SAC assemblies detected in the shuffled dataset (false positives or stochastic groups fortuitously appearing). Bottom, The estimated false positive rate (eFPR) for different alpha levels. eFPR was defined as the number of SAC assemblies detected on the shuffled dataset divided by the number of SAC assemblies detected in the original dataset. The alpha level selected in this manuscript (Z = 2.57) has an eFPR smaller than 0.25%. (B) eFPR for different numbers of cells within a dataset. (**C**) eFPR for the four selected alpha levels highlighted in B (dotted lines): Z=1.96 (yellow), 2.57 (red), 2.81 (blue), 3.29 (black), corresponding to p-values of 0.05, 0.01, 0.005, and 0.001, respectively. Curves show mean *±* s.e.m. across datasets. Because eFPR values differ substantially across alpha levels, the same data are shown at multiple y-axis zoom levels to allow visualization of both large-and small-amplitude variations. For all alpha levels, eFPR is constant for datasets with more than 40 cells.

**Supplementary Figure 5:**
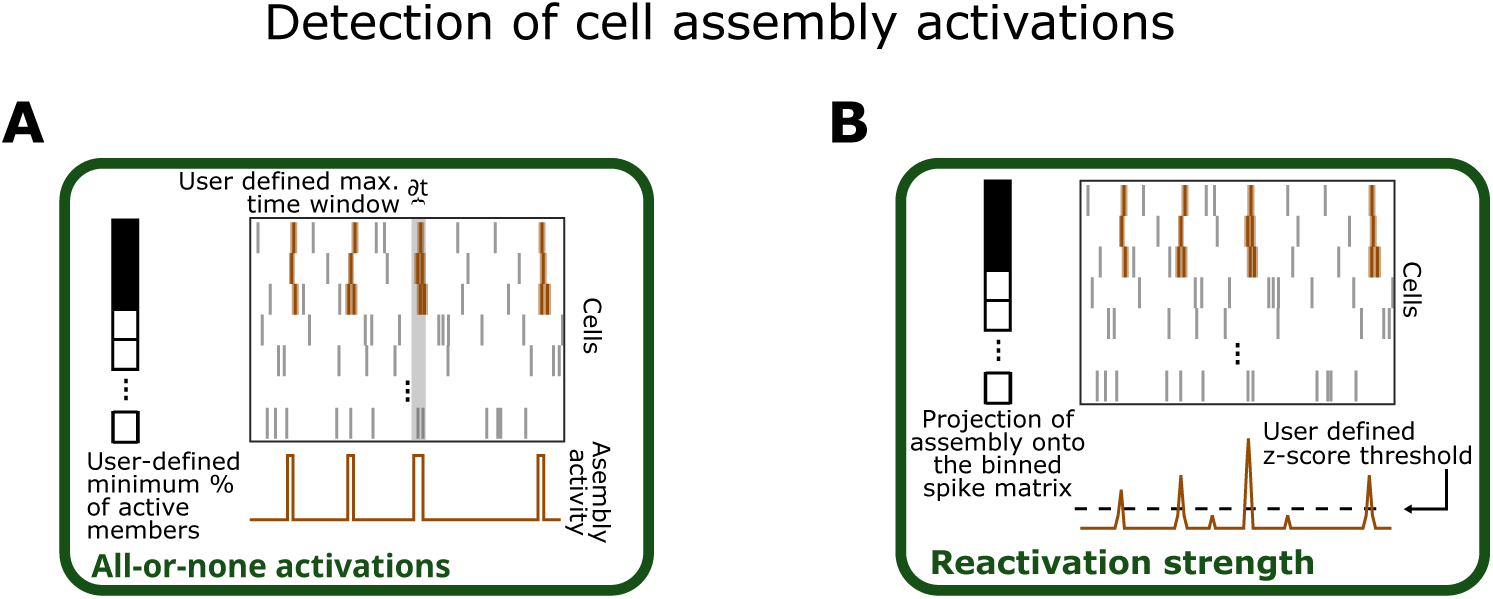
Methods for the detection of assembly activations. (A) Assembly activity is analyzed using an all-or-none binary approach. The user specifies the maximum temporal window (i.e., timescale) and the minimum proportion of assembly members required to be active (100% for strict analyses). Activations are defined by periods that satisfy both conditions. (B) Assembly assembly activity quantified as reactivation strength by projecting assembly composition vectors onto a binarized spike matrix. Assemblies are considered active when this activation measure exceeds a user-defined threshold (usually z-scored).

**Supplementary Figure 6:**
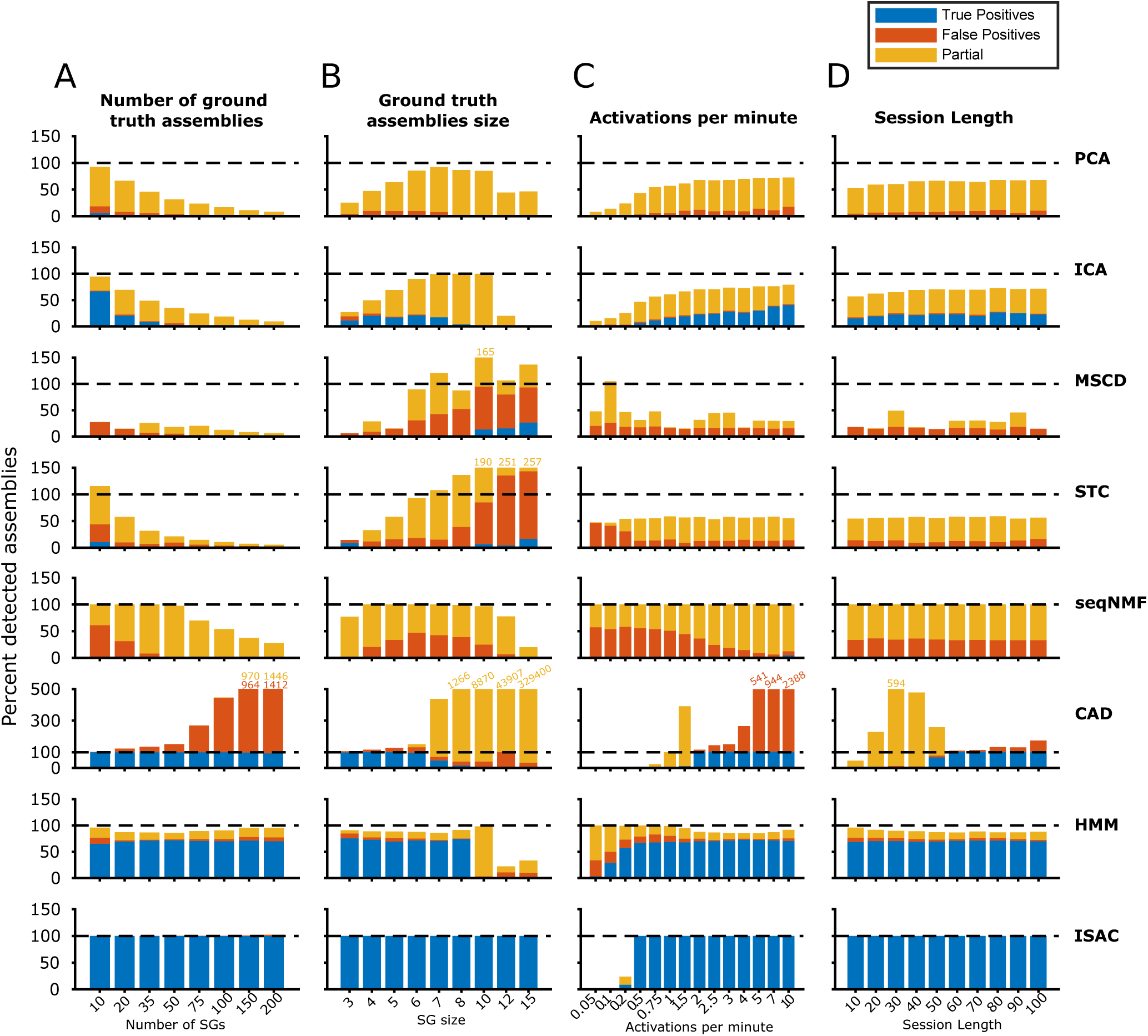
Detailed performance results on simulated datasets with Poisson point process as background noise (related to. Figure 4). Performance results shown as a percentage of the total number of ground truth assemblies. In same cases bars are beyond y scale limits and numbers above bars represent bar height (yellow number for yellow bar height, red number for red bar height).

**Supplementary Figure 7:**
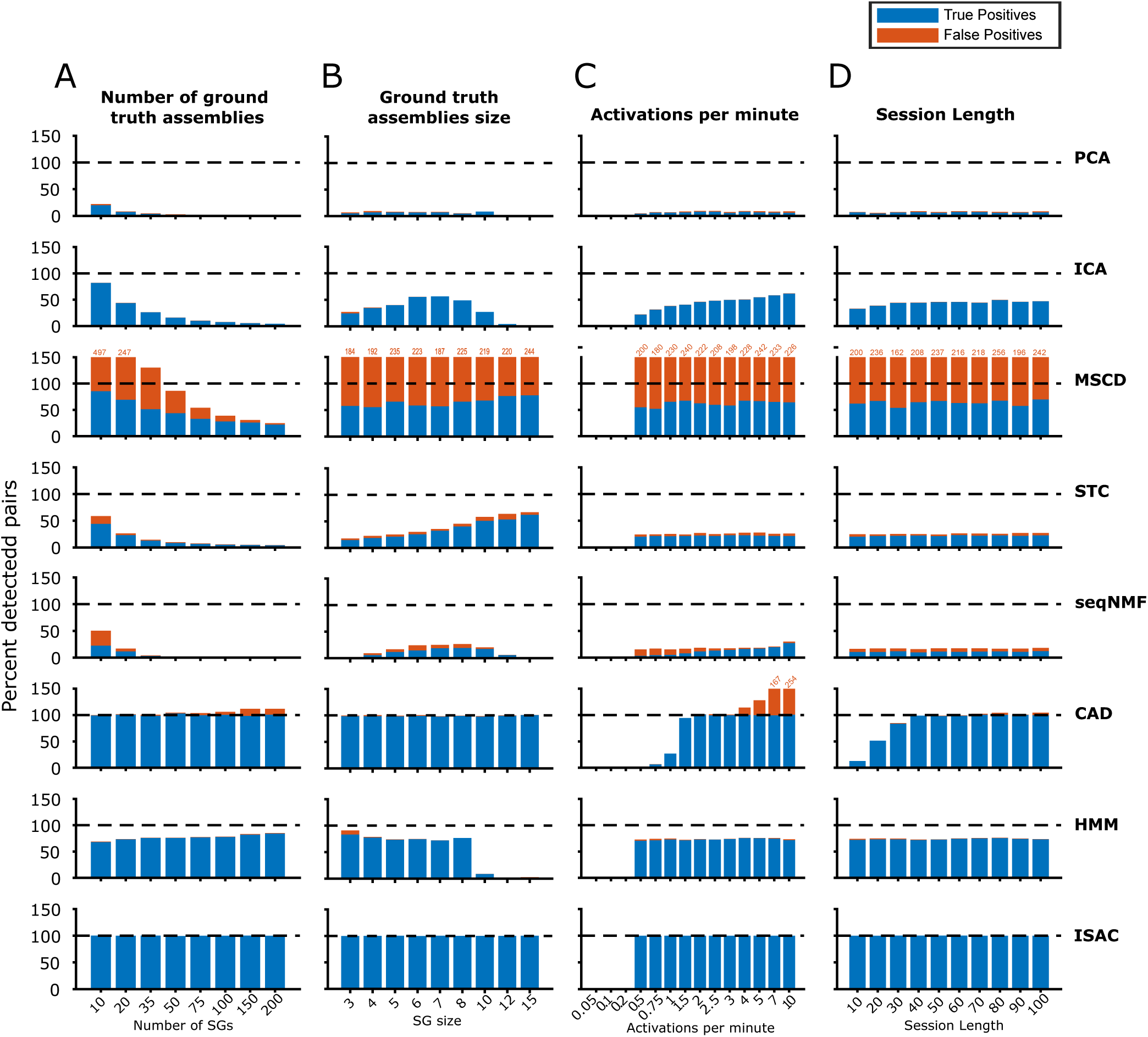
Algorithm performances in detecting pairs of synchronous cells within simulated recordings of ground truth assemblies embedded in Poisson point process as randomized background activity (related to. Figure 4). Performance is expressed as the percentage of correctly detected pairs relative to the total number of ground-truth pairs. Numbers above bars represent bar height (red number for red bar height).

**Supplementary Figure 8:**
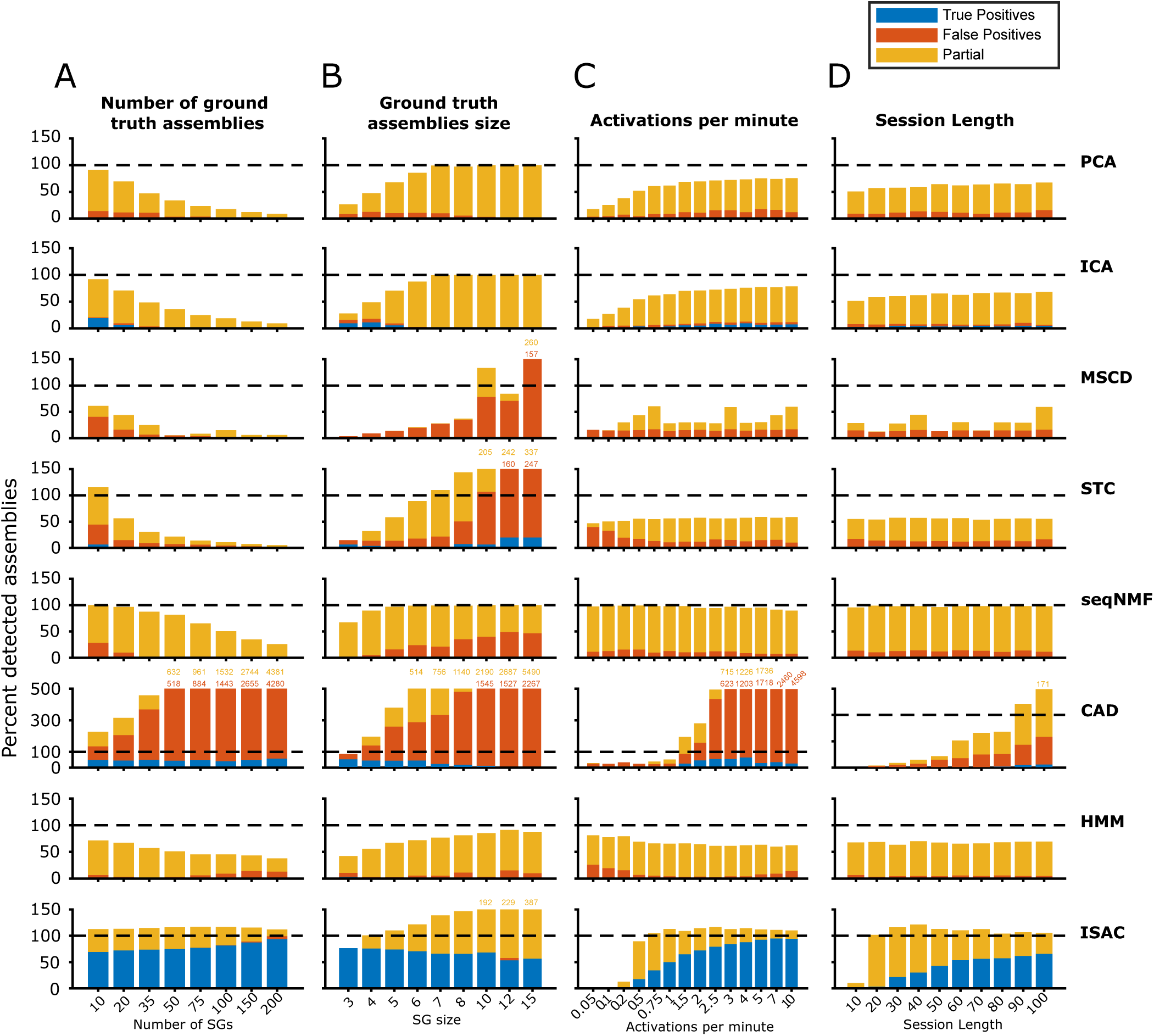
Detailed performance results on simulated datasets with shuffled prefrontal cortex recording as randomized background activity (related to. Figure 5). Same as Supp Figure 6 but related to Figure 5.

**Supplementary Figure 9:**
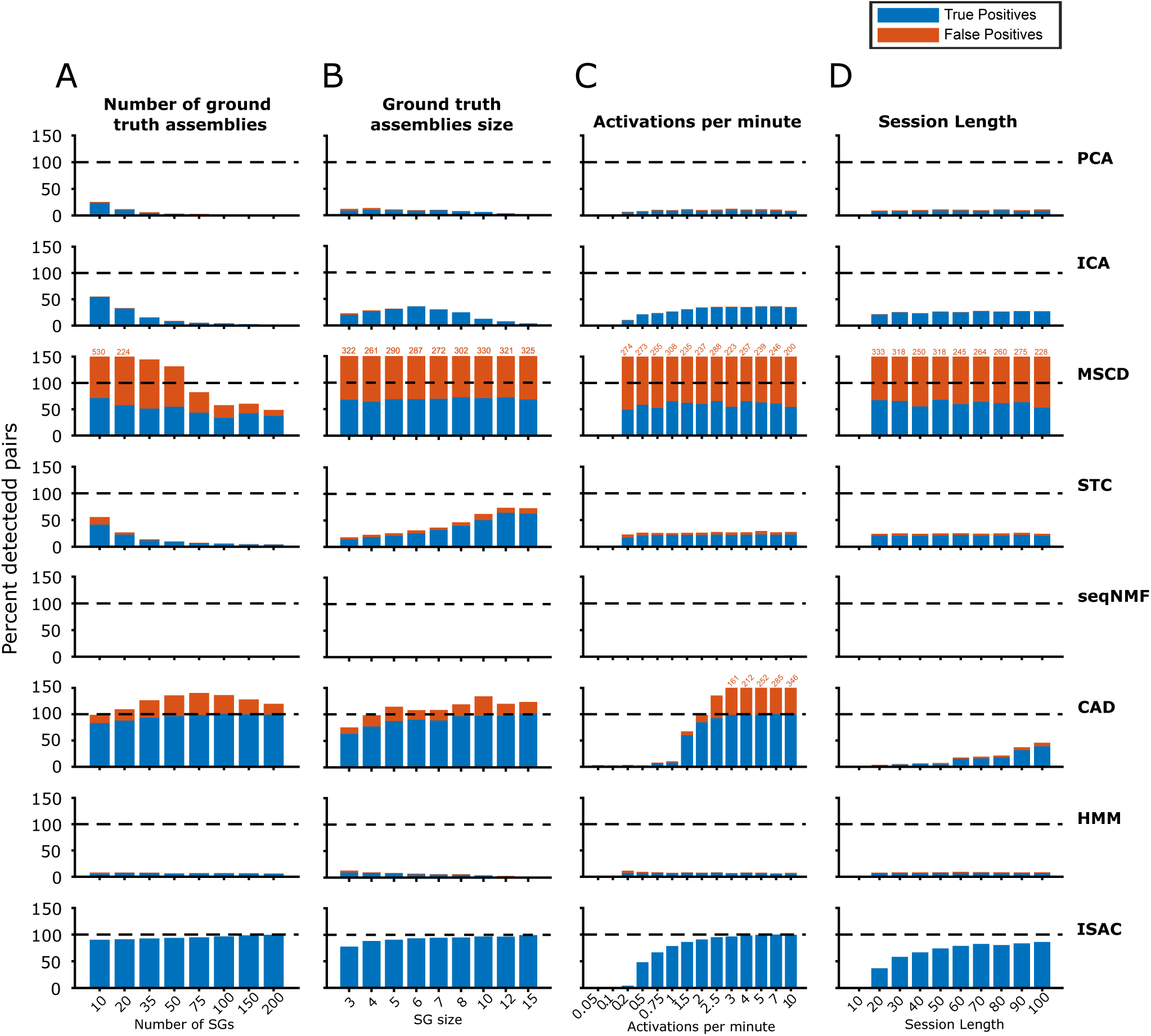
Algorithm performances in detecting pairs of synchronous cells within simulated recordings with shuffled prefrontal cortex recording as randomized background activity (related to. Figure 5). Performance shown as a percentage of the total number of ground truth pairs. Numbers above bars represent bar height (red number for red bar height).

**Supplementary Figure 10:**
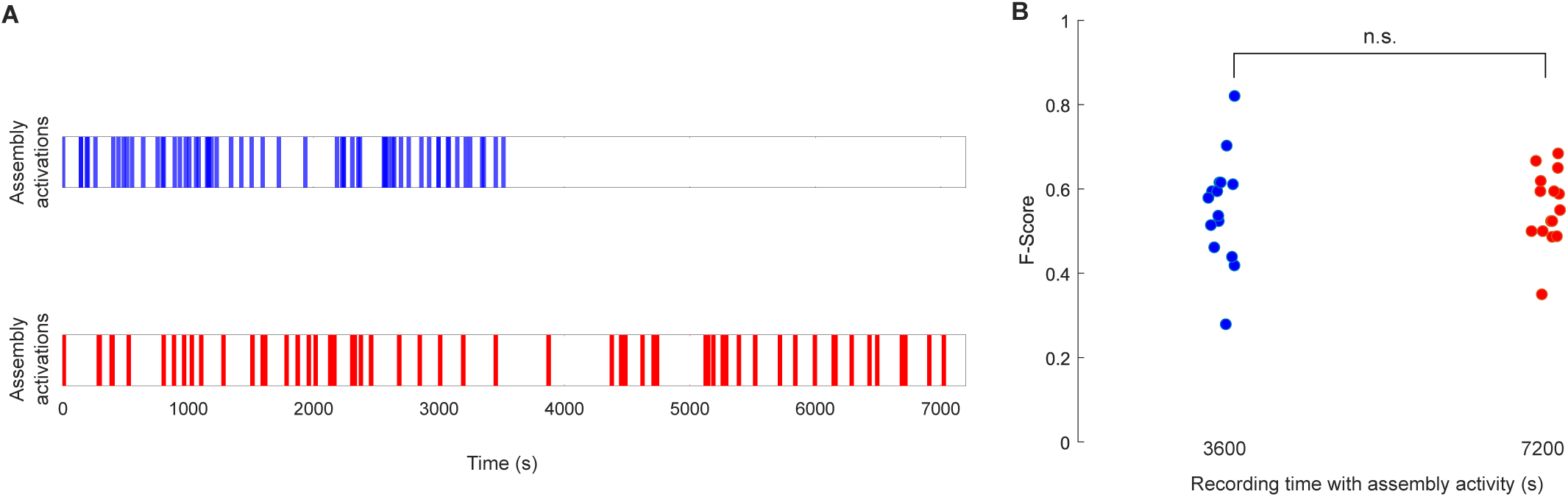
ISAC performance is preserved when assembly activity is temporally restricted. Recording time with assembly activity (s) (A) Example of assembly activity in two dataset types: one in which assembly activations are restricted to half of the recording duration, and one in whih assembly activations occur throughout the entire recording. (B) Comparison of ISAC performance in detecting assemblies across 15 datasets of each type shown in (**A**).

**Supplementary Figure 11:**
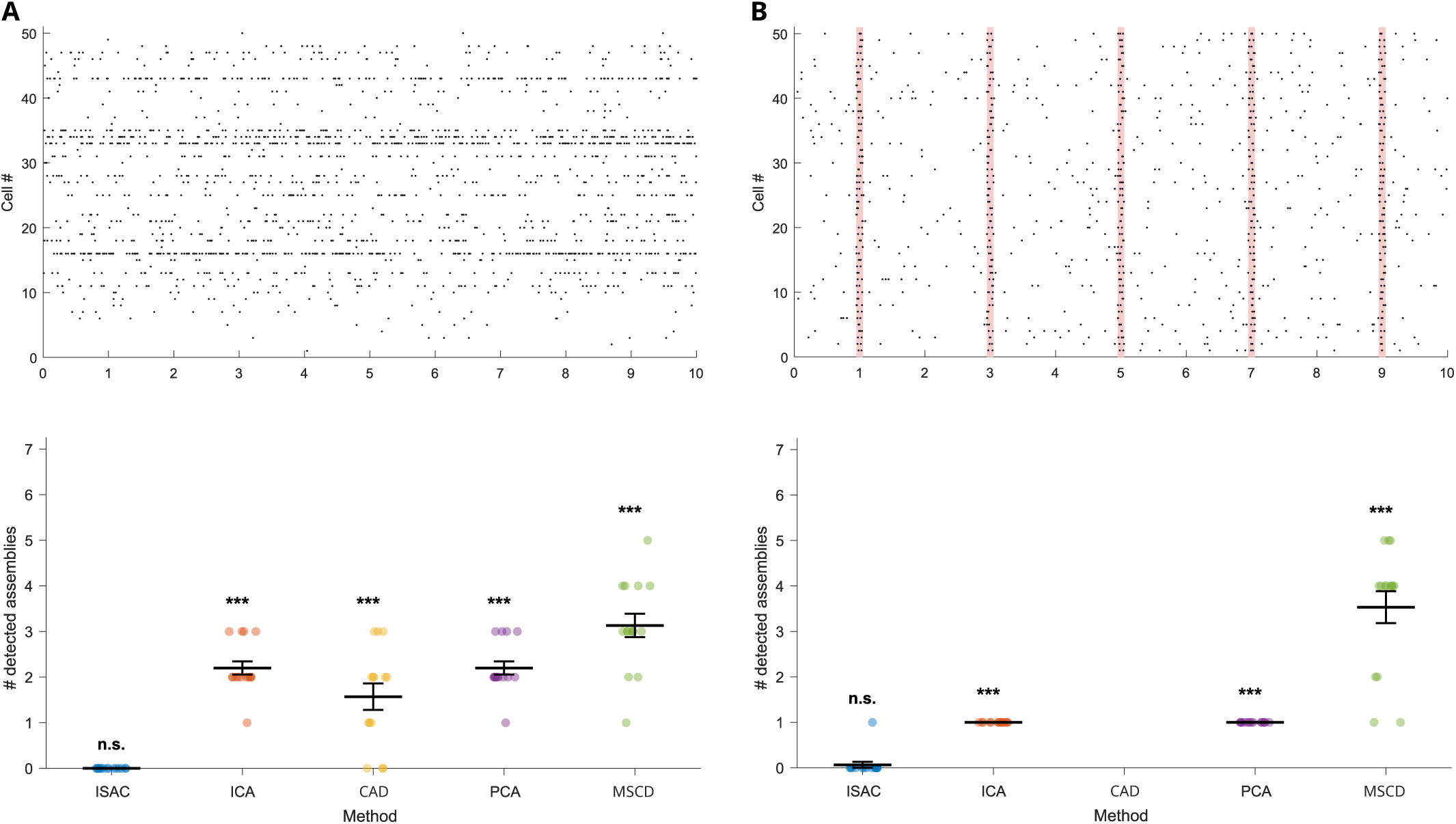
Absence of ISAC false-positive detections in datasets without ground-truth SAC assemblies. (A) Top, representative 10 s snapshot of the raster plot from a 1 h dataset without injected ground-truth assemblies, generated by shuffling real PFC recordings. Bottom, number of falsely detected cell assemblies identified by ISAC and the other methods across 15 datasets of this type. Stars indicate statistically significant differences from zero, whereas n.s. denotes a non-significant difference (Wilcoxon signed-rank test). HMM and seqNMF were not included in these tests because both methods require the number of assemblies to be specified in advance. Informing them that the dataset contained zero assemblies would have provided ground-truth information and therefore would not have constituted a fair comparison. (B) Top, same as in (**A**) but for a 1 h dataset without ground-truth assemblies and with Poisson background activity containing non-stationary events (red bands). Bottom, same as in (**A**) but across 15 datasets of this type. CAD was unable to complete a single run on these datasets, even after many hours of computation.

**Supplementary Figure 12:**
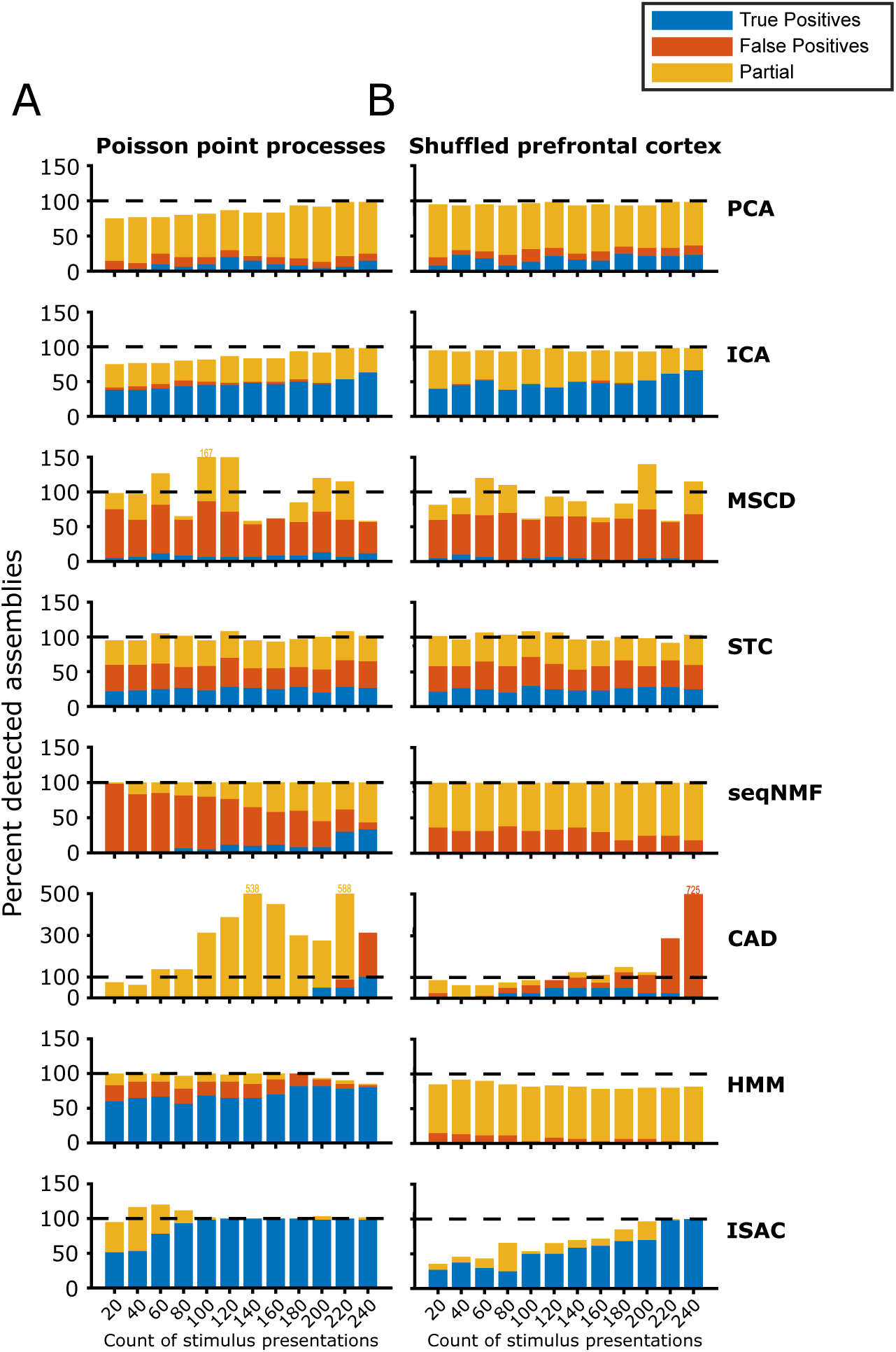
Detailed performance results in detecting SAC assemblies within simulated recordings of an auditory processing experiment (related to. Figure 6). (A) Performances in detecting SAC assemblies with Poisson point process as randomized background activity. (B) Performances in detecting SAC assemblies with shuffled prefrontal cortex recordings as randomized background activity. Numbers above bars represent bar height (yellow number for yellow bar height, red number for red bar height).

**Supplementary Figure 13:**
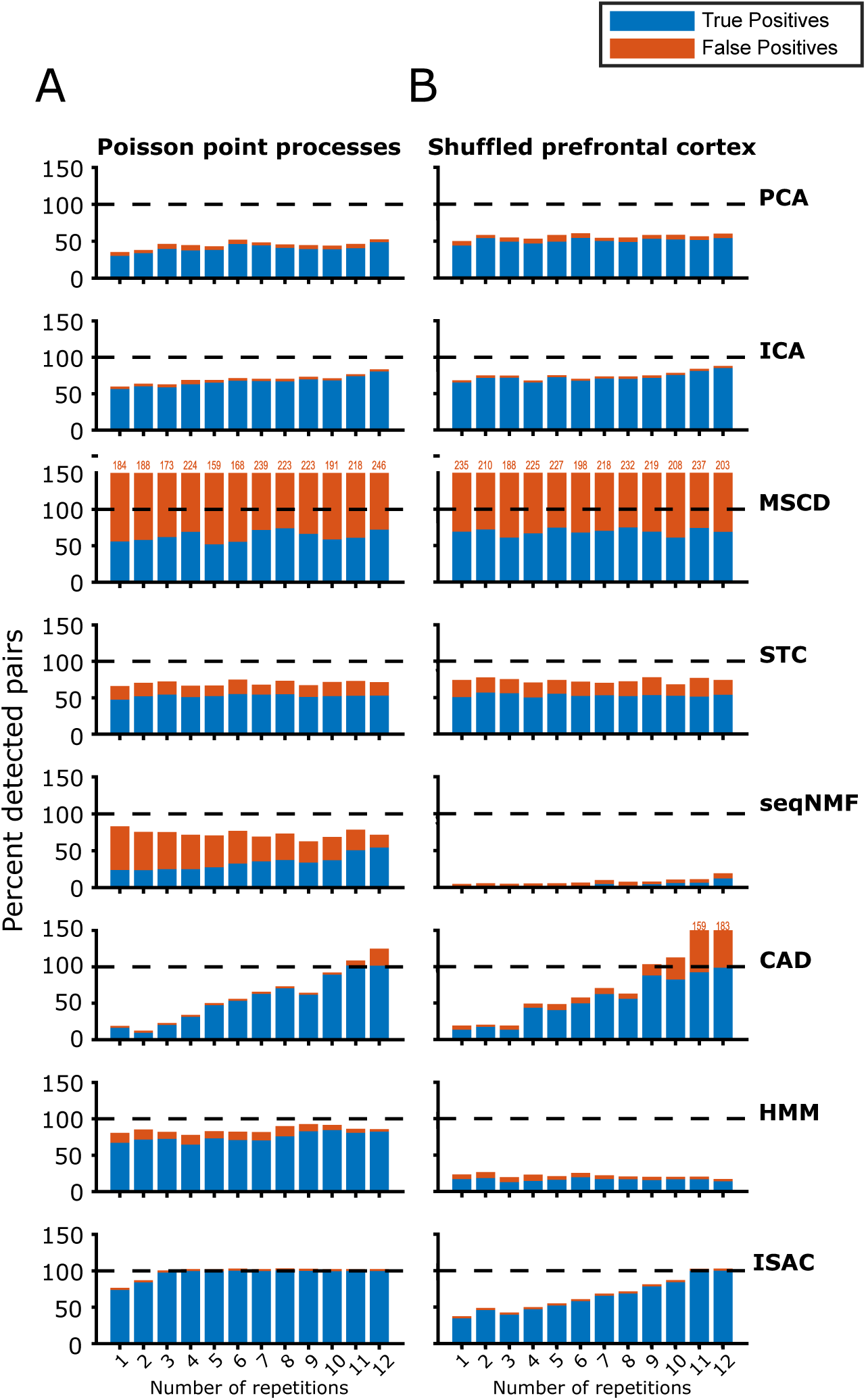
Performance results in detecting pairs of synchronous cells within simulated recordings of an auditory processing experiment (related to. Figure 6). (A) Performances in detecting pairs of synchronous cells with Poisson point process as background noise. (B) Performances in detecting pairs of synchronous cells with shuffled prefrontal cortex recordings as background noise. Numbers above bars represent bar height (red number for red bar height).

**Supplementary Figure 14:**
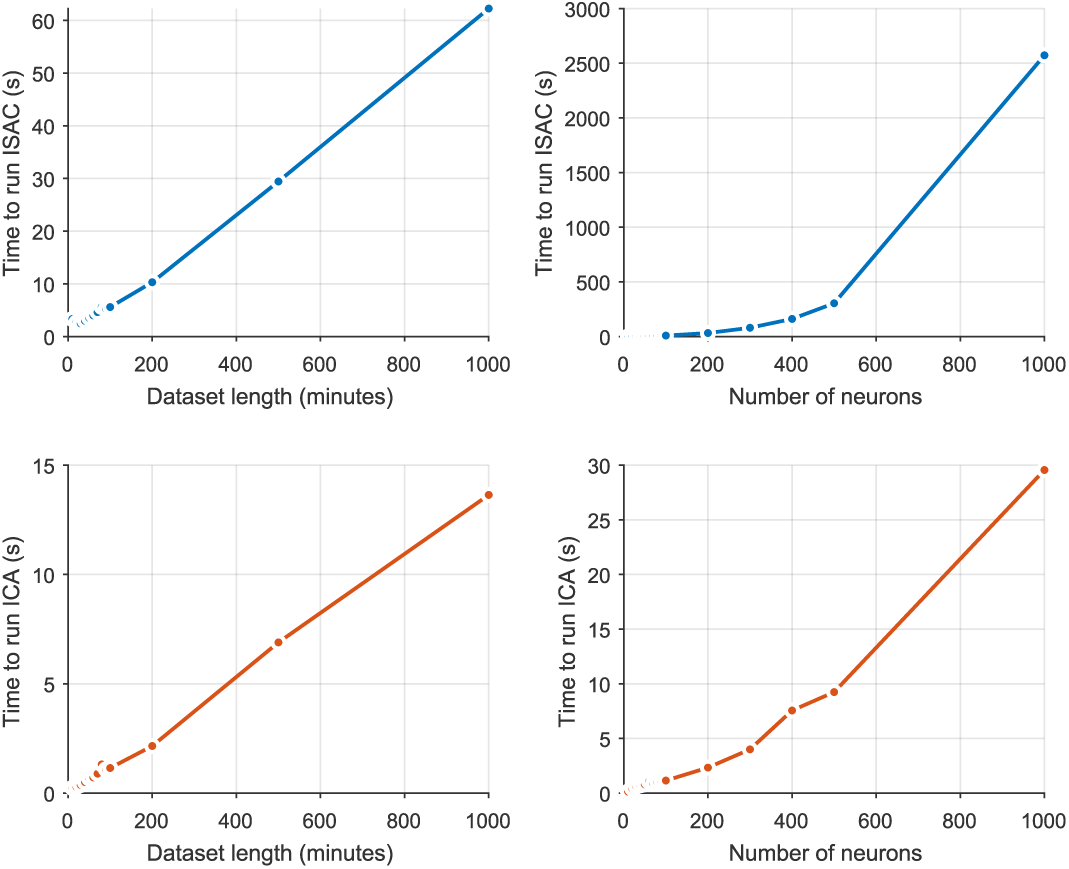
Computational cost of ISAC vs. ICA. Runtime as a function of recording duration (left) and number of recorded neurons (right) for ISAC (top) and ICA (bottom).

**Supplementary Figure 15:**
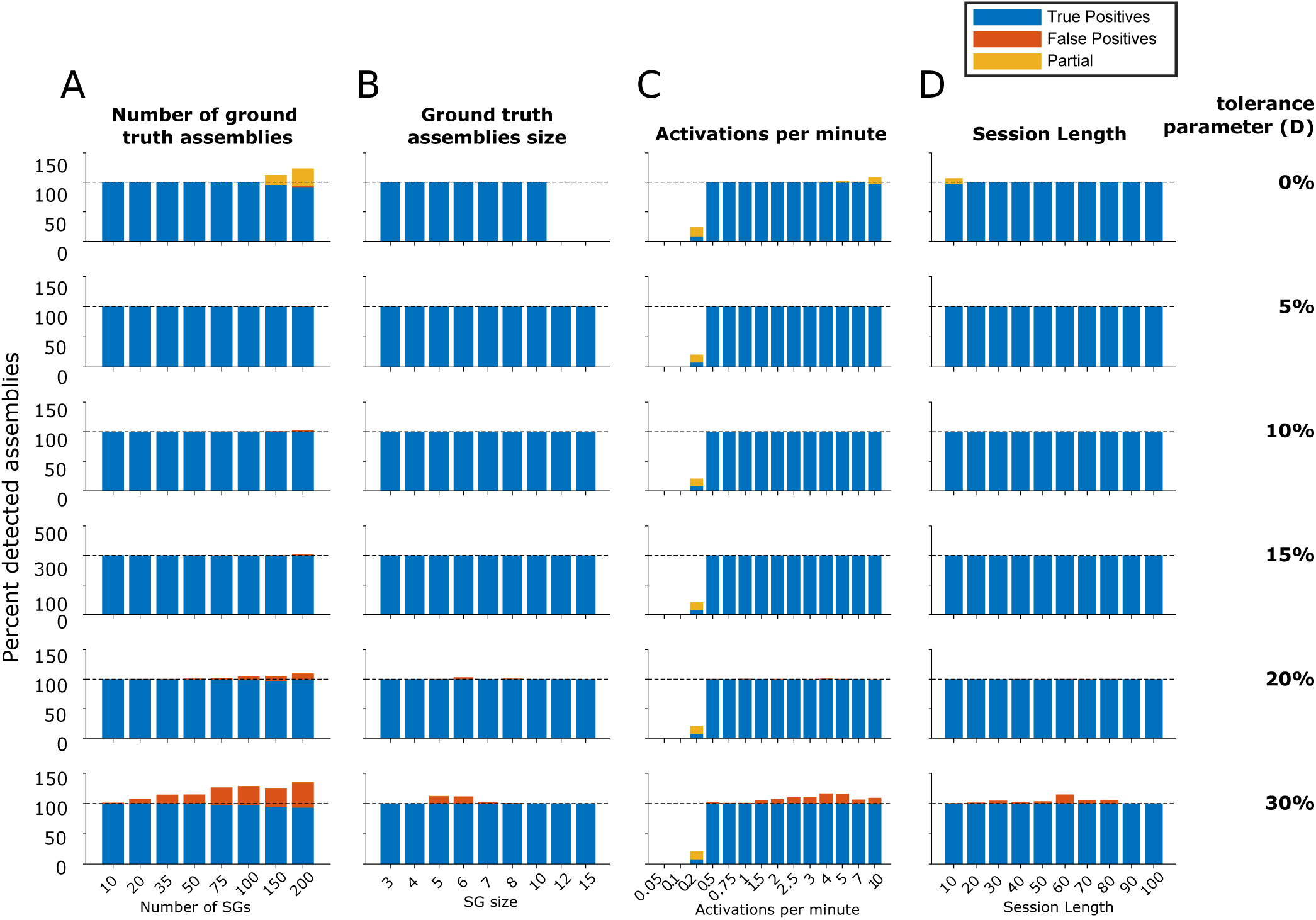
Detailed performance results on simulated datasets (Poisson point process as background noise) of ISAC using different values for the tolerance parameter, **D**. Results are shown as a percentage of the total number of ground truth assemblies. Numbers above bars represent bar height (yellow number for yellow bar height, red number for red bar height).

**Supplementary Figure 16:**
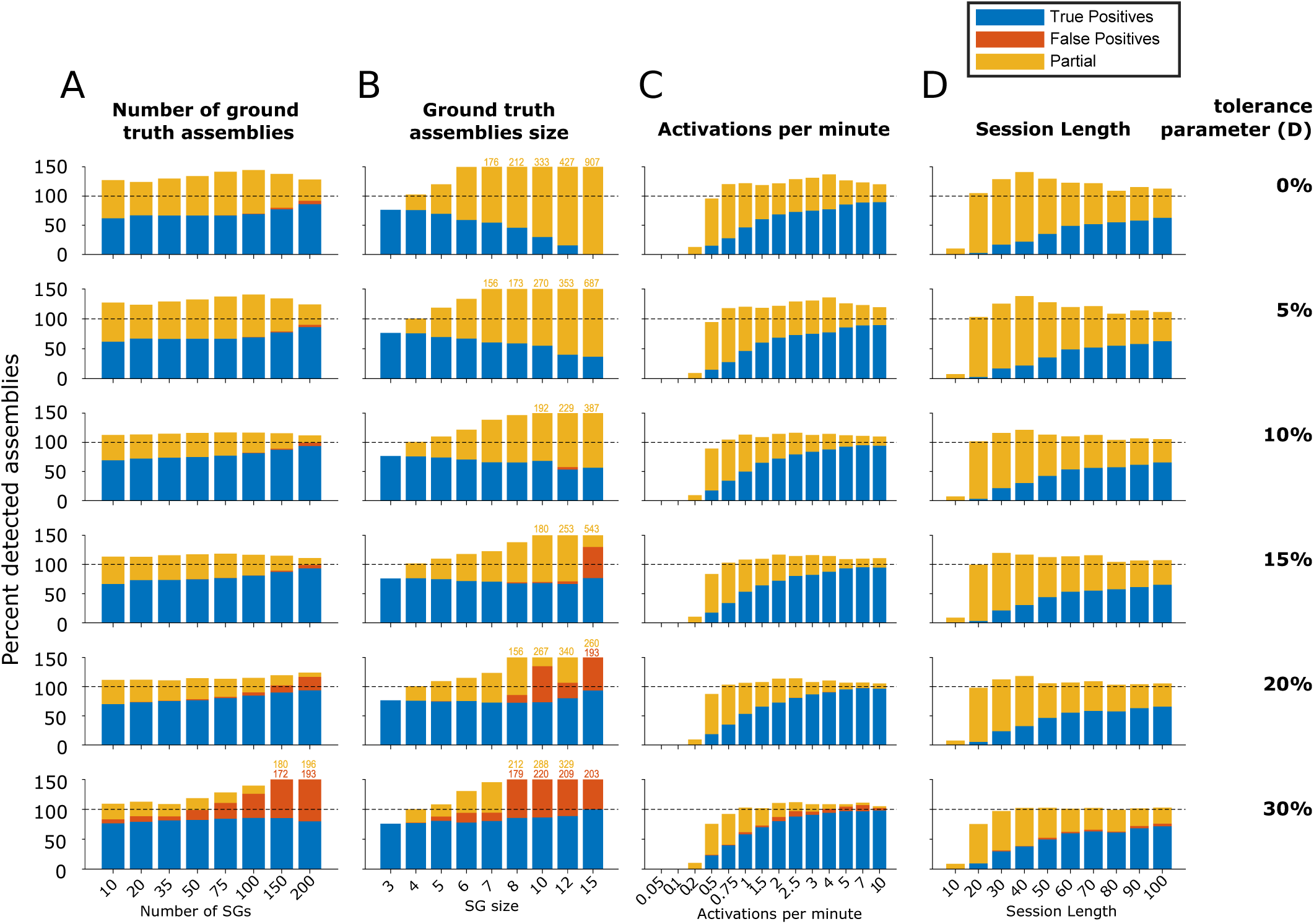
**Detailed performance results on simulated datasets (shuffled prefrontal cortex recording as background noise) of ISAC using different tolerance values**. Same as Supp. Figure 15 but relative to Figure 5.

**Supplementary table 1.** Characteristics of simulations. Simulations 42 to 53 correspond to Figure 6.

| Simulation | Number of SAC assemblies | Size of SAC assemblies | Session Length (min) | Number of activations |
| --- | --- | --- | --- | --- |
| 1 | 10 | 5 | 60 | 120 |
| 2 | 20 | 5 | 60 | 120 |
| 3 | 35 | 5 | 60 | 120 |
| 4 | 50 | 5 | 60 | 120 |
| 5 | 75 | 5 | 60 | 120 |
| 6 | 100 | 5 | 60 | 120 |
| 7 | 150 | 5 | 60 | 120 |
| 8 | 200 | 5 | 60 | 120 |
| 9 | 67 | 3 | 60 | 120 |
| 10 | 33 | 4 | 60 | 120 |
| 11 | 20 | 5 | 60 | 120 |
| 12 | 13 | 6 | 60 | 120 |
| 13 | 10 | 7 | 60 | 120 |
| 14 | 7 | 8 | 60 | 120 |
| 15 | 4 | 10 | 60 | 120 |
| 16 | 3 | 12 | 60 | 120 |
| 17 | 2 | 15 | 60 | 120 |
| 18 | 20 | 5 | 10 | 20 |
| 19 | 20 | 5 | 20 | 40 |
| 20 | 20 | 5 | 30 | 60 |
| 21 | 20 | 5 | 40 | 80 |
| 22 | 20 | 5 | 50 | 100 |
| 23 | 20 | 5 | 60 | 120 |
| 24 | 20 | 5 | 70 | 140 |
| 25 | 20 | 5 | 80 | 160 |
| 26 | 20 | 5 | 90 | 180 |
| 27 | 20 | 5 | 100 | 200 |
| 28 | 20 | 5 | 60 | 3 |
| 29 | 20 | 5 | 60 | 6 |
| 30 | 20 | 5 | 60 | 12 |
| 31 | 20 | 5 | 60 | 30 |
| 32 | 20 | 5 | 60 | 45 |
| 33 | 20 | 5 | 60 | 60 |
| 34 | 20 | 5 | 60 | 90 |
| 35 | 20 | 5 | 60 | 120 |
| 36 | 20 | 5 | 60 | 150 |
| 37 | 20 | 5 | 60 | 180 |
| 38 | 20 | 5 | 60 | 240 |
| 39 | 20 | 5 | 60 | 300 |
| 40 | 20 | 5 | 60 | 420 |
| 41 | 20 | 5 | 60 | 600 |
| 42 | 4 | 5 | 10 | 10 |
| 43 | 4 | 5 | 10 | 20 |
| 44 | 4 | 5 | 10 | 30 |
| 45 | 4 | 5 | 10 | 40 |
| 46 | 4 | 5 | 10 | 50 |
| 47 | 4 | 5 | 10 | 60 |
| 48 | 4 | 5 | 10 | 70 |
| 49 | 4 | 5 | 10 | 80 |
| 50 | 4 | 5 | 10 | 90 |
| 51 | 4 | 5 | 10 | 100 |
| 52 | 4 | 5 | 10 | 110 |
| 53 | 4 | 5 | 10 | 120 |

**Supplementary Table 2.** Parameters used for each benchmarked method. Ground-truth information was provided only when required by the original implementation. In our simulations, assembly spikes were distributed within a 30-ms jitter window. All methods require specification of a timescale over which cell assemblies are detected, and this timescale was therefore set to 30 ms for all methods requiring this parameter, with the exception of MSCD, which instead uses a biophysically inspired exponential kernel (*τ* = 3 ms), corresponding to a decay of approximately 15 ms and thus compatible with the 30-ms assembly timescale simulated here (assemblies at substantially different timescales are not detected under this parameterization).

| Method | Parameters used | Source code | Ground-truth info provided |
| --- | --- | --- | --- |
| **PCA**<br>(Peyrache et al. 2010) | 30 ms bins, 50% overlap<br>components retained above the Marchenko–Pastur bound | FMAToolbox<br>http://fmatoolbox.sourceforge.net | — |
| **STC**<br>(Humphries 2011) | 1 ms quantisation<br>15 ms Gaussian SD ($30\text{ms}/2$)<br>cosine similarity | Author's repository<br>https://github.com/mdhumphries/SpikeTrainCommunitiesToolBox | — |
| **ICA**<br>(Lopes-dos-Santos et al. 2013) | As PCA, then FastICA on the significant PCs subspace<br>Bin size = 30 ms | FMAToolbox<br>http://fmatoolbox.sourceforge.net | — |
| **MSCD**<br>(Billeh et al. 2014) | 1 ms bins<br>3 ms exponential constant<br>81 log-spaced Markov times (0.01–100)<br>optimal Markov time is the longest plateau in the Markov stability analysis as described by the authors | Author's repository<br>https://github.com/CellAssembly/Detection | — |
| **CAD**<br>(Russo & Durstewitz 2017) | 30 ms bins<br>max lag 2<br>$\alpha = 0.05$ (default) | Author's repository<br>https://github.com/DurstewitzLab/Cell-Assembly-Detection | — |
| **HMM**<br>(Maboudi et al. 2018) | 30 ms bins<br>number of assemblies<br>PBEs at MUA > 3 SD as described by the authors | Author's repository<br>https://github.com/nelpy | number of assemblies in simulation |
| **seqNMF**<br>(Mackevicius et al. 2019) | 30 ms bins<br>factor length 1<br>$K$ = number of assemblies | Author's repository.<br>https://github.com/FeeLab/seqNMF | number of assemblies in simulation |
| **ISAC** | $\Delta t = 30$ ms<br>$Z = 2.57$<br>$D = 10\%$ | Here<br>https://github.com/mnpompili/ISAC | — |

